# Metals and metal isotopes in insect wings: Implications for diet, geolocation and pollution exposure

**DOI:** 10.1101/2022.11.02.514901

**Authors:** Megan S. Reich, Mira Kindra, Felipe Dargent, Lihai Hu, D.T. Tyler Flockhart, D. Ryan Norris, Heather Kharouba, Gerard Talavera, Clément P. Bataille

**Affiliations:** Department of Biology, University of Ottawa, Ottawa, ON, Canada; Department of Earth and Environmental Sciences, University of Ottawa, Ottawa, ON, Canada; Great Lakes Forestry Centre, Canadian Forest Service, Natural Resources Canada, Sault Ste. Marie, ON, Canada; Department of Integrative Biology, University of Guelph, Guelph, ON, Canada; Institut Botànic de Barcelona (IBB), CSIC-Ajuntament de Barcelona, Barcelona, Catalonia, Spain

**Keywords:** Metal isotopes_1_, strontium isotope ratios_2_, lead isotopes_3_, isotope-based geographic assignment_4_, chemoprints_5_, monarch butterfly_6_, metal pollution_7_

## Abstract

Anthropogenic activities are exposing insects to abnormal levels of toxic metals, with unknown implications for migratory insects. Simultaneously, metals and metal isotopes have become promising tools for the geolocation of migratory insects. Furthering our understanding of metal cycling in insect tissues is essential, both for the development of metals and metal isotopes as geolocation tools, and for assessing the toxicity risks of metals to insects. We conducted a diet-switching experiment on monarch butterflies (*Danaus plexippus*) with controlled larval and adult diets to evaluate the dietary and environmental sources of 23 metals and metalloids, strontium isotopes, and lead isotopes to insect wing tissues over a period of 8 weeks. Concentrations of Ca, Co, and Sb differed between the sexes. Ni and Zn bioaccumulated in the insect wing tissues over time, likely from the adult diet, while increases in Al, Cr, Cd, Cu, Fe, and Pb were likely from external sources (i.e., dust aerosols). Bioaccumulation of Pb in the monarch wings was confirmed by Pb isotopes to be from external anthropogenic sources, revealing the potential of Pb isotopes to become an indicator and tracer of metal pollution exposure along migratory paths. Concentrations of Ba, Cs, Mg, Na, Rb, Sr, Ti, Tl, and U appeared to be unaffected by dietary or environmental contamination and should be further developed for geolocation purposes. Strontium isotope ratios remained indicative of the larval diet, at least in males, supporting its potential as a geolocation tool. However, the difference in strontium isotope ratios between sexes, as well as the possibility of external contamination by wetting, requires further investigation. Our results demonstrate the complexity of metal cycling in insects and the need for further investigations, as well as the value of studying metals to develop new tools to quantify pollution exposure, metal toxicity and insect mobility.

## 1 Introduction

Out of the 20 elements considered essential for human life, ten are metals. Although this list can differ among taxa, several metals are essential to the survival and functioning of organisms. It is estimated that up to 30% of proteins are metalloproteins (Aptekmann *et al*. 2022), and essential metals participate in enzyme catalysis, protein structure, cellular signaling, and the creation of electrochemical potentials at membranes (Maret 2016). However, essential metals follow a dose-response curve, in which low doses cause morbidity due to metal deficiency and high doses cause toxicological effects. Conversely, many non-essential metals are toxic even at low doses, although some show hormesis (i.e., a beneficial effect at low doses of an otherwise toxic substance). Therefore, maintaining metal homeostasis is important for survival, and organisms have evolved mechanisms for the uptake, transport, sequestration, and removal of metals. Despite the fundamental importance of metals, little is known about the pathways of metal cycling and incorporation in insects (Dow 2017).

Understanding the sources and mechanisms of metal incorporation into insect tissues is especially important in the context of anthropogenic habitat alteration because of the adverse effects that human-induced changes in metal concentrations can have on insect populations. The rapid, worldwide declines of many insect populations has key implications for ecosystem health, conservation, and biosecurity, and has led to widespread calls to conservation action (Dirzo *et al*. 2014). Anthropogenic influences are thought to be the main cause of population declines via the effects of climate change, habitat loss and degradation, increased use of pesticides, and the resultant changes in biotic interactions, such as phenological mismatches (Wagner 2020). Furthermore, activities such as mining, industrialization, and agriculture, are causing insects to be exposed to metals at rates above the natural baselines they evolved to tolerate (Monchanin *et al*. 2021). Even when regulated, these activities can have adverse effects on insects because invertebrates are more susceptible than vertebrates to toxic metals like cadmium and lead and, therefore, often experience adverse effects well below anthropocentric permissible concentration limits (Hopkin 1990, Mogren and Trumble 2010, Monchanin *et al*. 2021). Migratory species could be at an even higher risk of adverse effects from metal exposure than non-migratory species (Seewagen 2020). Migration is a complex behavior, and migratory insects can travel hundreds to thousands of kilometers and are potentially exposed to many sources of metal pollution along their migratory routes. However, the potential effects of metals on migratory insect survival and migratory behavior are virtually unknown. To understand the conservation and toxicity risks of metal to insects, particularly migratory insects, we must first understand how metals are transported from the environment to insect tissues.

Besides these toxicological concerns, the analysis of metal and metal isotopes in insect tissues has become a promising avenue for developing tools aimed at understanding insect migration. Metals (e.g., Tigar and Hursthouse 2016), metal ratios (e.g., Holder *et al*. 2014), and combined metal composition (i.e., chemoprints, mineral component analysis, chemometrics; e.g., Lin *et al*. 2019) have been used to assess long-term patterns of insect dispersal. Most recently, metal isotopes have been applied to trace the natal origin of migratory insects (Holder *et al*. 2014, Reich *et al*. 2021). These metal biomarkers are particularly valuable for tracking migratory insects because traditional biologging techniques are usually unviable for insects due to (1) their large and rapidly fluctuating populations, (2) their relatively short lifespans and multi-generational migratory cycles, and (3) the dynamic spatiotemporal connectivity of their populations (Garcia-Berro *et al*. 2022, López-Mañas *et al*. 2022). The development of these geolocation techniques requires that the given metals or metal isotopes vary predictably over the landscape such that baseline maps (i.e., “metalscapes” or “isoscapes”) can be developed as a reference for insect tissues of unknown geographic origin. Metals and metal isotopes are incorporated into insect tissue from the environment during tissue formation and give the resulting tissue a distinctive metal or metal isotopic signature that reflects the local environment in which the tissue was grown (e.g., Flockhart *et al*. 2015). Ultimately, the metal or metal isotopic signature of insect tissue is compared to the metal-specific baseline maps to estimate the location where tissue formation occurred (i.e., the natal origin), a process known as geographic assignment. However, making proper inferences with these tools requires a clear understanding of metal and metal isotope cycling in insect tissue over time because alterations to the natal signature, for example, via uptake of metals through adult feeding, will decrease the accuracy of geographic assignments (Dempster *et al*. 1986).

It is assumed that metals and metal isotopes, such as strontium isotope ratios, analyzed in insect wings (i.e., the preferred tissue for isotope geolocation studies in insects) come primarily from the diet and remain unchanged after wing formation (e.g., Holder *et al*. 2014, Lin *et al*. 2021, Reich *et al*. 2021). However, the potential addition of metals to insect wings after tissue formation has not been explicitly investigated. Recent controlled studies on keratin, metabolically inert tissues often used in the isotope geolocation of vertebrates (e.g., feathers: Crowley *et al*. 2021, hair: Fauberteau *et al*. 2021), demonstrated that some metals in keratin are rapidly replaced by exogenous contamination sources (e.g., wet absorption, pollution, dust aerosols; Hu *et al*. 2018, 2019, Rodiouchkina *et al*. 2022). Additionally, unlike hair, insect wings are living tissue primarily composed of chitin, a structural polysaccharide, and proteins. Insect wings host androconial organs (i.e., pheromone glands), sensory receptors, trachea, nerves, and hemolymph circulate through their veins (Tsai *et al*. 2020). Insect hemolymph transports nutrients and wastes, and provides a link between the wings and the insect body, thus providing an additional pathway for the potential ‘contamination’ of natal signatures, as has been observed for hydrogen isotope values (Lindroos *et al*. this issue). Thus, metals in insect wings are likely initially incorporated from the larval diet, but this initial endogenous signal could be altered by both internal (e.g., dietary) and external (e.g., dust aerosols) sources, depending on the properties and cycling of the specific metal.

In this study, we aimed to better characterize the cycling and incorporation mechanisms of a suite of metal and metal isotopes in insect wings over time and advance the development of metal-based tools for insect ecology and toxicology. We conducted a laboratory-based diet-switching experiment to explore the environmental and dietary sources of 23 metals and metalloids, strontium isotopes (measured as ^87^Sr/^86^Sr ratios), and lead isotopes (measured as ^208^Pb/^206^Pb and ^207^Pb/^206^Pb ratios) in insect wings over an 8-week timespan. For each metal, we evaluated the effects of intrinsic factors on metal concentrations and assessed the potential sources of any bioaccumulation. We also assessed the potential for ^87^Sr/^86^Sr in insect wings to be altered by wetting. Finally, we evaluated the potential of the metal and metal isotopes for providing dietary, geolocation, or pollution exposure information. By understanding the cycling of metals and metal isotopes in insect wings over time, we will be better able to test the assumptions of geolocation using metal biomarkers and assess the metal exposure risks of insects.

## 2 Materials and Methods

We chose for this study a representative winged insect, the monarch butterfly (*Danaus plexippus* (L.)). The monarch butterfly is an iconic migratory insect famous for its annual, multi-generational, round-trip migration from the Oyamel fir forests of Central Mexico, through the USA, to Canada, and back. Monarchs are a well-suited species to investigate the cycling, incorporation, and homeostasis of metal and metal isotope signals. First, they have a large geographic range with known elemental and isotopic variations meaningful for geolocation (Wieringa *et al*. 2020, Reich *et al*. 2021). Second, monarch larvae are specialist herbivores dependent on milkweed (*Asclepias* spp.). Plants are known to uptake and regulate metals in a species-specific manner; thus, this relatively high host-specificity will likely minimize potential discrepancies in metal signatures associated with differences in larval host plant species (McLean *et al*. 1983, Tibbett *et al*. 2021). Third, metal exposure is of particular concern for monarchs because milkweed is often found in disturbed habitats, such as roadsides, which often correlates with higher pollution exposure (Phillips *et al*. 2021). As for most insects, the cycling and toxicology of metals in monarch butterflies are mostly unknown, although zinc has been shown to decrease larval survival (Shephard *et al*. 2020).

To explore the cycling of metal and metal isotopes, we first conducted a diet-switching experiment on monarch butterflies over an 8-week period. We raised larvae from eggs with controlled milkweed dietary inputs and then maintained the adults with a controlled adult diet. We measured the changes in metal concentrations and isotopes in monarch wings at weekly time intervals and compared the metal abundances found in our laboratory experiment to the range of natural metal abundances in the wings of 100 wild-caught monarch butterflies. Next, we explored the incorporation mechanisms of different metals by calculating enrichment factors to the adult diet and a proxy of mineral dust for each metal, analyzing lead isotopes in the monarch wings to assess the source of Pb bioaccumulation, and performing a wetting experiment for Sr. Finally, we measured the concentrations of 12 metals in ashed milkweed samples from across the eastern USA and assessed our ability to use soil and milkweed samples in the construction of “metalscapes” for monarch butterflies (Supplementary Material).

### 2.1 Diet-switching experiment

We conducted a diet-switching experiment in the summer of 2019 to assess the sources and preservation of metals and metalloids (hereinafter referred to collectively as metals) and metal isotope signals in monarch wings throughout the life of an adult butterfly. Eggs were collected from a single wild-caught female, allowing us to assume that all the specimens in this experiment are at least half-siblings and control for maternal effects. After collection, eggs were transferred to individual 500 mL glass rearing containers with a mesh lid and placed in a controlled environmental chamber (Biochambers Inc., model LTCB-19) held at a light to dark schedule of 12h: 12h, light temperature of 28°C and dark temperature of 26°C, and humidity of 60%. These conditions reflect optimal conditions for monarch development (Dargent *et al*. 2021).

Caterpillars were fed wild common milkweed (*Asclepias syriaca* L.) leaves that were collected daily from August 9 to August 22, 2019, from a single 30 m^2^ site in Ottawa, Ontario, Canada (45.41°, −75.67°). The collection site is in an urban park next to the Rideau River and is near a light rail station and bike paths. The leaves were cleaned of surface contaminates (e.g., dust) using distilled deionized (DDI) water before being fed *ad libitum* to the caterpillars. Sub-samples of the cleaned milkweed leaves were collected for analysis to check the assumption that milkweed leaves taken from the same locality will have similar elemental and isotopic signatures over the 2-week period of collection. Frass was removed from the containers daily.

After the monarchs eclosed from the chrysalises (Aug. 29-31, 2019), they were transferred to mesh butterfly cages in an indoor laboratory. Butterflies were kept two per cage, but a cloth divider was installed to minimize wing damage from collisions between individuals. The cages were misted twice a day with DDI water to maintain humidity and were given a light-to-dark schedule of about 12h: 12h. The temperature in the laboratory was maintained by the building’s normal operations and maintenance schedule at approximately 20°C. Once per day, the adult butterflies were given 50% maple syrup solution in a tube with a foam faux-flower perch and allowed to feed *ad libitum*. Because the butterflies do not always recognize the unnatural food source, individuals were also hand-fed the maple syrup for at least one minute every other day. The first six monarchs were sampled immediately post-eclosion (Day 0; no feeding), and two monarchs were subsequently sampled each week until up to eight weeks post-eclosion (n = 22). Immediately after sampling, monarchs were placed into a freezing chamber at −18°C. Later, wings were dissected from the body and placed in glassine envelopes until elemental analysis. The time between collection and elemental or isotopic analysis ranged from a few months to 18 months.

#### 2.1.1 Trace element analysis

Elemental analysis of monarch wings, larval diet (i.e., milkweed leaves) and adult diet (i.e., 50% maple syrup solution) was performed in two analytical runs via inductively coupled plasma mass spectrometry (ICP-MS; Agilent 8800 ICP-QQQ, Agilent Technologies Inc., Santa Clara, CA, USA) at the Department of Earth and Environmental Sciences, University of Ottawa. The first analytical run in Feb. 2020 measured 23 metals in monarch wings (n = 18) and maple syrup (n = 4): aluminium (Al), arsenic (As), barium (Ba), calcium (Ca), cadmium (Cd), cobalt (Co), chromium (Cr), copper (Cu), caesium (Cs), iron (Fe), magnesium (Mg), manganese (Mn), molybdenum (Mo), sodium (Na), nickel (Ni), lead (Pb), rubidium (Rb), antimony (Sb), strontium (Sr), titanium (Ti), thallium (Tl), uranium (U), and zinc (Zn). To remove any surface contaminants (e.g. dust), a single forewing from each monarch was cleaned using pressurized nitrogen gas for about 2 min. at 10 psi (Reich et al. 2021). The monarch wings and maple syrup samples were then digested in perfluoroalkoxy alkanes (PFA) vials (Savillex, LLC., Eden Prairie, MN, USA) using 1 mL 16 M HNO_3_ (double distilled TraceMetal™ Grade; Fisher Chemical, Mississauga, ON, Canada) and 0.1 mL ultraclean hydrogen peroxide (for trace analysis Grade; Sigma-Aldrich, Italy) for 12 hrs. at 120 °C. After digestion, the samples were dried on a hot plate at 90 °C and then re-dissolved in 1 mL of 2% v/v HNO_3_ (double distilled). A 100 μL aliquot of each solution was extracted, diluted to 2 mL 2% v/v HNO_3_, and centrifuged. Calibration standards were prepared using single element certified standards (SCP Science Inc., Montreal, QC, Canada). Detection limits were conservatively estimated as three times the standard deviation of the blanks within each run. For monarch wings, 4 measurements of As, 2 of Tl, and 8 of U were below the detection limit. Concentrations of Cd, Cs, Tl, and U were lower than the calibration range of 1.56 to 100 ppb (~200, ~800, ~900, and ~4600 times lower, respectively). Metal concentrations in maple syrup were comparatively low, with all 4 As and U measures and 1 Fe measure below the detection limit. Additionally, Cd, Tl, and Pb were all well below the calibration range (~600, ~200 and ~200 times lower, respectively). Intra-run relative standard deviation (RSD) of the repeated standard analysis was below 3% except for Cs (RSD = 21%), Sb (RSD = 19%), Rb (RSD = 67%), and Fe (RSD = 38%).

For the second analytical run in May 2021, milkweed samples (n = 3) and analytical replicates of monarchs wings (n = 19) were analyzed for 15 metals: Al, Ba, Ca, Cd, Co, Cr, Cu, Fe, Mg, Mn, Ni, Pb, Sr, Ti, and Zn. Milkweed samples were cleaned with DDI water, an ultrasonic bath (10 min.), and a MilliQ water rinse. Wing samples were dry-cleaned with pressurized nitrogen gas for 10 min. at about 10 psi for a limited, repeated elemental analysis of the monarch wings to assess if there was sample contamination of Pb during storage. For monarch tissues, a single hindwing was used for most samples. However, some samples had too low Pb concentrations and both hindwings (i.e., L2) or both hindwings and the remaining forewings were used (i.e., L11, L16, L17, L21, L5). Samples were then digested in concentrated nitric acid (16 M; distilled TraceMetal™ Grade; Fisher Chemical, Canada) using microwave digestion (Anton Paar Multiwave 7000, Austria): the samples were heated from ambient temperature to 250 °C at a steady rate in 20 min. and then left at 250 °C for 15 min. in a pressurized chamber. An aliquot of 200 μL from each sample was separated and diluted to 2 mL of 2% v/v HNO_3_ (double distilled) and run on the ICP-MS. For monarch wings, 16 measurements of Co and 17 of Cd were below the detection limit; only Cd was lower than the calibration range (~155 times lower). Metal concentrations in milkweed were comparatively high, with Ca and Mg well above the calibration range (~900 and ~200 times higher, respectively). Intra-run RSD of the repeated standard analysis was below 3% except for Mg (RSD = 70%), Ca (RSD = 157%), and Fe (RSD = 143%).

#### 2.1.2 Strontium isotope ratios analyses

The ^87^Sr/^86^Sr analysis was performed in two runs at the Isotope Geochemistry and Geochronology Research Centre, Carleton University using a ThermoScientific™ Neptune™ high-resolution multi-collector inductively coupled plasma mass spectrometer (MC-ICP-MS; Thermo Fisher Scientific, Bremen, Germany). To prepare for the first run in Feb. 2020, milkweed samples (n = 3) were cleaned using DDI water and an ultrasonic bath (10 min.), then oven-dried at 80 °C before being ashed at 600°C for 4 hrs. to facilitate digestion. They were then digested in perfluoroalkoxy alkanes (PFA) vials (Savillex, LLC., Eden Prairie, MN, USA) using 1 mL 16 M HNO_3_ (double distilled TraceMetal™ Grade; Fisher Chemical, Mississauga, ON, Canada) and 0.1 mL ultraclean hydrogen peroxide (for trace analysis Grade; Sigma-Aldrich, Italy) for 12 hrs. at 120 °C. After digestion, the samples were dried on a hot plate at 90 °C. Dried samples were re-dissolved in 1 mL of 2% v/v HNO_3_ (double distilled). A 100 μL aliquot of each solution was extracted, diluted to 2 mL 2% v/v HNO_3_, and the concentration of Sr was measured using the ICP-MS in Oct. 2019. The remaining ~900 μL aliquot (90%) of sample digest was dried down and re-dissolved in 0.05 mL 6 M HNO_3_. The separation of Sr was processed in 100 μL microcolumn loaded with Sr-spec Resin™ (100–150 μm; Eichrom Technologies, LLC). The matrix was rinsed out using 6 M HNO_3,_ and Sr was collected with 0.05 M HNO_3_. After separation, eluates were dried and re-dissolved in 2 mL 2% v/v HNO_3_ for ^87^Sr/^86^Sr analysis.

For the second analytical run in Sept. 2020, the remaining aliquots (~90%) of the monarch wing (n = 18) and maple syrup (n = 4) samples run on the ICP-MS in Feb. 2020 were used. Samples were dried down and re-dissolved in 0.05 mL 6 M HNO_3_. The separation of Sr was processed in 100 μL microcolumn loaded with Sr-spec Resin™ (100–150 μm; Eichrom Technologies, LLC). The matrix was rinsed out using 6 M HNO_3,_ and Sr was collected with 0.05 M HNO_3_. The microcolumn process was repeated to ensure complete separation. After separation, the eluates were dried, and monarch wing samples were re-dissolved in 200 μL 2% v/v HNO_3_ and maple syrup in 2 mL 2% v/v HNO_3_ for ^87^Sr/^86^Sr analysis.

For both analytical runs, sample solutions were introduced to the MC-ICP-MS using a micro*FAST* MC single-loop system (Elemental Scientific Inc., Omaha, NE, USA). The sample introduction flow rate was 30 μL/min, and the loading volume was 200 μL for the milkweed and maple syrup and 100 μL for the monarch wing samples. The solution was introduced using a PFA nebulizer, double-pass quartz spray chamber, quartz torch, and nickel sample and skimmer cones. Isotopes ^82^Kr, ^83^Kr, ^84^Sr, ^85^Rb, ^86^Sr, ^87^Sr, and ^88^Sr were simultaneously measured in L4, L3, L2, L1, C, H1, and H2 Faraday cups, respectively. Measurements of samples were made using a static multi-collector routine that consisted of one block of either 70 (milkweed and maple syrup) or 30 cycles (monarch wings) with an integration time of 4.194 s/cycle. ^84^Sr and ^86^Sr have isobaric interferences from ^84^Kr and ^86^Kr, respectively. ^87^Sr has an isobaric interference from ^87^Rb. The interferences of ^84^Sr and ^86^Sr were corrected by subtracting the amount of ^84^Kr and ^86^Kr corresponding to the ^83^Kr signal. Interference of ^87^Sr was corrected by subtracting the amount of ^87^Rb corresponding to the ^85^Rb signal. Instrumental mass fractionation was corrected by normalizing ^86^Sr/^88^Sr to 0.1194 using the exponential law (Moore *et al*. 1982). Strontium isotope compositions are reported as ^87^Sr/^86^Sr ratios. Procedural blanks were always negligible, with good reproducibility of ^87^Sr/^86^Sr for NIST SRM987 (0.710251 ± 0.000003 (1 SD), n = 2). Two 100 ng/g pure Sr standards were measured along with the samples as in-house standards (SrSCP: 0.70816 ± 0.00013, n = 11; GSC: 0.70756 ± 0.00002, n = 4). The long-term reproducibility of the SrSCP in-house standard is (0.70822 ± 0.00004, n = 106).

#### 2.1.3 Lead isotope ratios analysis

We analyzed the lead isotope ratios (^208^Pb/^206^Pb, ^207^Pb/^206^Pb) of monarch wing and milkweed samples to refine the sources of Pb in the monarch wings. The remaining aliquots (~90%) of the microwave-digested milkweed (n = 3) and monarch (n = 19) samples run on the ICP-MS in May 2021 were used. The separation of Pb was processed in polyethylene columns (Bio-Rad, Hercules, California, USA) loaded with 0.6 mL analytical grade anion exchange resin (AG 1-X8, 100-200 mesh, chloride form; Bio-Rad, Hercules, California, USA). Initial rinsing steps used 1 M HBr (OPTIMA™, Fisher Chemical, Canada) followed by the collection of the Pb fraction with ultra-clean 6 M HCl. A second pass through the column was performed to further purify Pb. After separation, the Pb eluates were dried and re-dissolved in 260 μL and 460 μL 2% v/v HNO_3_, for monarch and milkweed samples, respectively (TraceMetal™ Grade; Fisher Chemical, Canada). A thallium spike (in a Tl:Pb mass ratio of 1:4) was added to each sample for mass bias correction against ^203^Tl/^205^Tl = 0.418922.

The Pb isotopes analysis was performed at the Isotope Geochemistry and Geochronology Research Centre, Carleton University using a Thermo Scientific™ Neptune™ high-resolution multi-collector inductively coupled plasma mass spectrometer (MC-ICP-MS; Thermo Fisher Scientific, Bremen, Germany). Sample solutions in 2% v/v HNO_3_ were aspirated using a PFA nebulizer, double-pass quartz spray chamber, quartz torch, and nickel sample and skimmer cones. Isotopes ^202^Hg, ^203^Tl, ^204^Pb, ^205^Tl, ^206^Pb, ^207^Pb, and ^208^Pb were simultaneously measured in L3, L2, L1, C, H1, H2 and H3 Faraday cups, respectively. Measurements of samples were made using a static multi-collector routine that consisted of 1 block of 70 cycles for milkweed and 35-40 cycles for monarchs with an integration time of 8.389s/cycle. Procedural blanks were negligible (<50 pg). The reported Pb isotope ratios are corrected for offsets between the analyzed and reported NBS981 (Todt *et al*. 1996). For a period of 12 months, average ratios and uncertainties (± SD, n = 47) of NBS981 bracketing samples were ^206^Pb/^204^Pb = 16.9310 ± 0.0095, ^207^Pb/^204^Pb = 15.4851 ± 0.0012, ^208^Pb/^204^Pb = 36.6783 ± 0.0035, ^208^Pb/^206^Pb = 2.16634 ± 0.00011, and ^207^Pb/^206^Pb = 0.91460 ± 0.00003.

#### 2.1.4 Statistical analysis

To examine patterns in metal concentrations and isotopes in the monarch wings over time, and to group elements and isotopes with similar characteristics, we used multivariate multiple regression and principal component analysis (PCA). General linear models were constructed with sex, time (days), and body mass (mg) as fixed effects; multiple models were run independently because there were differences in sample size due to exclusions from the quality control process. All models were visually checked for normality, independence, and homoscedasticity of the model residuals. Body mass and sex are strongly associated (point-biserial correlation = −0.70; Figure S3), so linear models were re-run without sex as an explanatory variable to test the sensitivity of the results to the confounding variables (Table S3). Twenty metals were included in the PCA because of unequal sample size due to As, Tl, and U having measurements below the detection limit. All statistical analyses were performed using R (version 4.1.0; R Core Team 2013), and a commented R markdown document has been provided (see Data Availability Statement).

#### 2.1.5 Mobility index and enrichment factors

To explore the transfer of metals between trophic levels, we calculated the elemental mobility index for each metal between the larval diet (i.e., milkweed) and the monarchs sampled on the day of eclosion (i.e., day 0). The mobility index was calculated as the ratio between the metal concentration in the monarch wings and the larval diet (Tigar and Hursthouse 2016). A value greater than one indicates that the monarch wing has a higher metal concentration than its larval food and that the metal is biomagnified; whereas a value of less than one indicates that the monarch wing has lower metal concentrations than its larval food and the metal has been biodiluted.

Enrichment factors (EF) were also calculated between the adult diet and monarch wings (EF_ms_) and between the upper continental crust (Rudnick and Gao 2014) and monarch wings (EF_cc_) to assess the source of metals in butterfly wings. The upper continental crust is used as a proxy for average exogenous contamination from mineral dust, which allows the EF to represent how much of a trace element is sourced endogenously (i.e., from within the organism itself) versus obtained through exogenous contamination (i.e., from the continental crust) (Hu et al. 2019). We calculated the EF using calcium concentration as a normalization factor (Kabata-Pendias & Mukherjee, 2007; Equation 1). Ca is one of the major elements in the continental crust and an abundant metal in monarch wings (Table S2); normalization to Ca makes the EF calculation less sensitive to variation between samples (Hu *et al*. 2019). An EF of greater than one implies that the metal is enriched in the monarch relative to the source (e.g., diet (EF_ms_), crust (EF_cc_)), and, therefore, the source has low potential to contaminate the monarch signal. Conversely, an EF of less than one indicates that this metal in monarch wings is susceptible to contamination from that source.

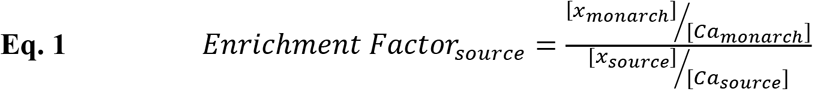

### 2.2 Wetting experiment

Because ^87^Sr/^86^Sr are currently the main metal isotopic tool used in insects, we conducted an additional experiment to refine the sources of Sr in wings and assess the potential contamination of Sr due to wetting. Submerging keratin (e.g., hair) in water can result in exogenous contamination of Sr (Hu *et al*. 2020), but this contamination route is untested for insect chitin. In our diet-switching experiment, the adult monarchs are not wetted, and, therefore, the Sr content and ^87^Sr/^86^Sr ratios are uncontaminated by wetting. However, in the wild, monarchs might be wetted by rain or other environmental waters, which could possibly lead to exogeneous Sr contamination via exchange, removal, or addition. Since butterfly wings are superhydrophobic (Pass 2018), we hypothesized that the Sr in the wings would not be contaminated by submergence in water. We designed a simple wetting experiment to test whether exogeneous Sr could be integrated into the monarch wings when submerged in water. Two beakers of 80 mL 300 ppb Sr certified standard purchased from SCP Science Inc. (Montreal, QC, Canada) were prepared and neutralized with 60 μL 0.5 N NaOH. Two monarch butterfly wings were cleaned of surface dust using pressured nitrogen gas (10 psi for 4 min.). The wings were submerged in the beakers of solution and agitated. At designated time steps, from before the wings were submerged to 28 days after submergence, a 100 μL aliquot of the solution was removed from each beaker, diluted to 3 mL 2% v/v HNO_3_ (TraceMetal™ Grade; Fisher Chemical, Canada), and centrifuged. These aliquots were then measured using the ICP-MS to monitor the change of Sr concentration in the beakers over time.

### 2.3 Natural metal concentrations in monarchs

To estimate the natural elemental concentration of metals in wild monarch butterflies, the wings of 100 monarch butterflies from eastern North America were analysed for a suite of 12 elements (i.e., Al, Ba, Cd, Co, Cr, Cu, Mg, Mn, Ni, Pb, Sr, and Zn). The butterflies were captured in the spring of 2011 from sites in Texas, Oklahoma, and Missouri as part of a previous study (Flockhart *et al*. 2013). Methodological details can be found in the Supplementary Material.

## 3 Results

### 3.1 Metals classified into four categories

The concentrations of Al, As, Ba, Ca, Cd, Co, Cr, Cs, Cu, Fe, Mg, Mn, Mo, Na, Ni, Pb, Rb, Sb, Sr, Ti, Tl, U, and Zn in the wings of the monarch butterflies, the larval diet (i.e., milkweed), and the adult diet (i.e., maple syrup) from the diet-switching experiment are reported in Table S2. In the monarch wings, Ca was the most abundant metal (mean ± SD; 640 ± 160 ng/mg, n = 18) followed by Mg (290 ± 180 ng/mg, n = 18) and Fe (87 ± 34 ng/mg, n = 18). Uranium was the least abundant (0.0010 ± 0.0003 ng/mg, n = 10), with many samples falling below the detection limit. In the larval diet, alkaline earth metals Ca, Mg, and Sr were the most abundant (Ca: 18 ± 2 μg/mg, n = 3; Mg: 4.2 ± 0.3 μg/mg, n = 3; Sr: 190 ± 20 ng/mg, n = 3) and Cd was the least abundant (9.5 ± 2.2 ng/g, n = 3). Similarly, in the adult diet Ca (190 ± 37 ng/mg, n = 4) and Mg (73 ± 5, ng/mg, n = 4) were the most abundant and Cd the least abundant (0.4 ± 0.1 ng/g, n = 4).

Principal component analysis (PCA) was used to help categorize the metals in the monarch wings from the diet-switching experiment. The first principal component (PC1) explained 43.8% of the variation and was driven mainly by Al, Fe, Pb, Cr, and Cs (Figures 1A and S1A). A linear regression of PC1 with time category (i.e., early, late) and sex (R^2^ = 0.80) detected a difference in PC1 between Week 0 and Weeks 1 to 8 (β = 5.5 ± 0.9, 6.3, p < 0.001; Figure 1B), but not between the sexes (p = 0.1). PC2, driven by Cu, Na, Mn, Ca, and Cd (Figure S1B), accounted for 16.1% of the variation. A linear regression with day and sex (R^2^ = 0.52) found that PC2 trended temporally (β = 0.05±0.02, 3.5, p = 0.003) and differed between the sexes (β = 2.3 ± 0.6, 3.7, p = 0.002), suggesting that the metals driving PC2 may be influenced by biological variables (Figure 1B).

**Figure 1.**
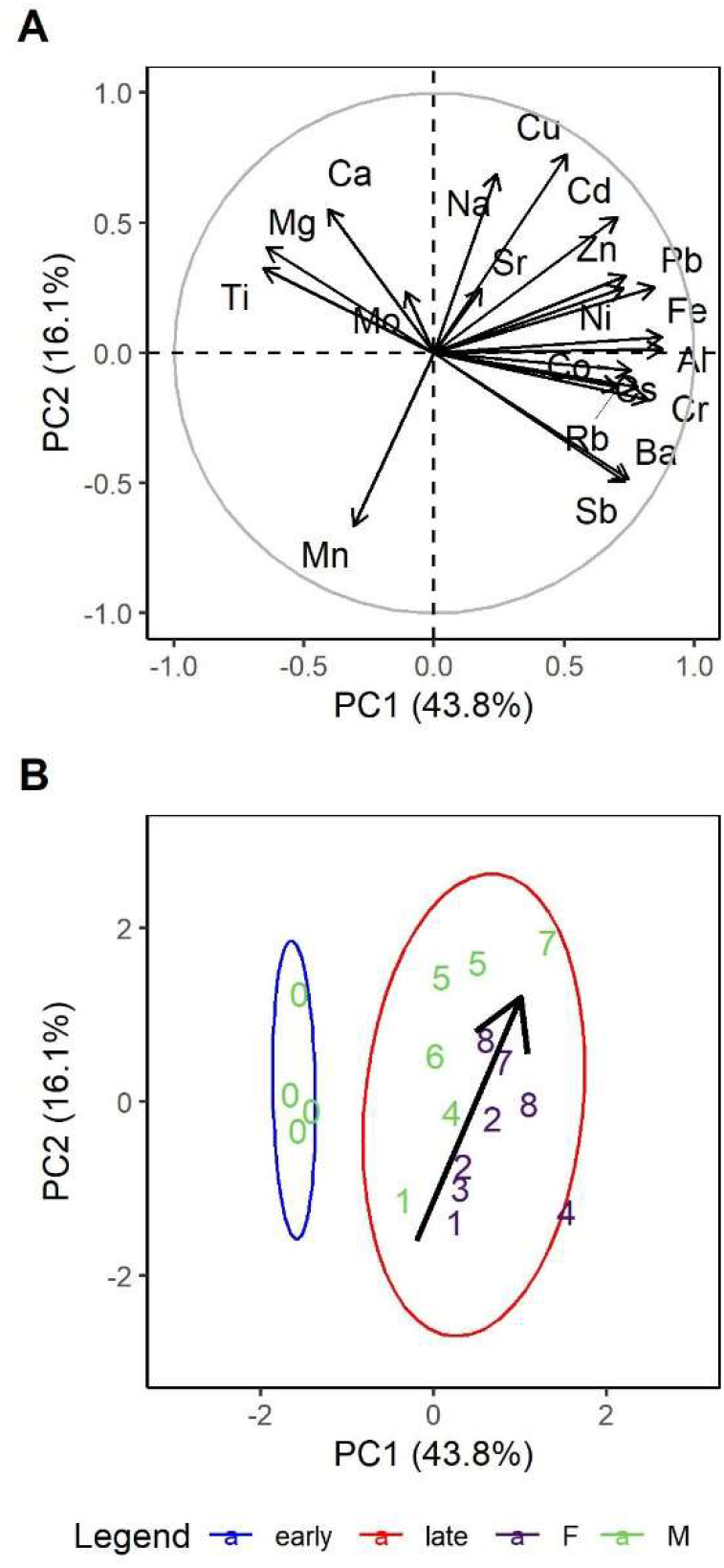
Principal components analysis biplots of twenty metals (excluding As, Tl, and U due to measurements below the detection limit) in monarch butterfly wings **(A)** Loading plot of the first two principal components for monarch wings from all weeks explaining 59.9% of the variation. **(B)** PCA score plot of monarch wings from all weeks; numbers depict the week at which the individual was sampled. There is a no overlap between the 95% data ellipses of early dates (i.e., Week 0; blue ellipse) and later dates (i.e., Weeks 1 to 8; red ellipse). There is a trend, mainly in PC2, from Week 1 to 8; a black arrow has been superimposed on the biplot to illustrate this trend. There is also grouping between female (purple) and male (green) butterflies.

Multivariate multiple regression found that metals in monarch wings showed distinctive associations with the time (days), sex, and body mass (mg; Table 1). If metals are accumulated from endogenous or exogenous sources, we would expect a positive correlation between time and metal concentration; such associations were found for Al, Cd, Cu, Fe, Pb, Ni, and Zn. Significant negative correlations with time were found for As and Mn, suggesting that these metals are removed from the butterfly wings over time. Calcium concentrations in male monarch wings were, on average (± SE), 190 ng/mg (± 60) higher than in females (Figure 2A). Conversely, on average males had 0.04 ng/mg (± 0.01) less Co and 1.0 ng/mg (± 0.5) less Sb. Body mass was found to have a significant association with Mo (β = −0.0006 ± 0.0002, −2.6, p < 0.05).

**Table 1.**
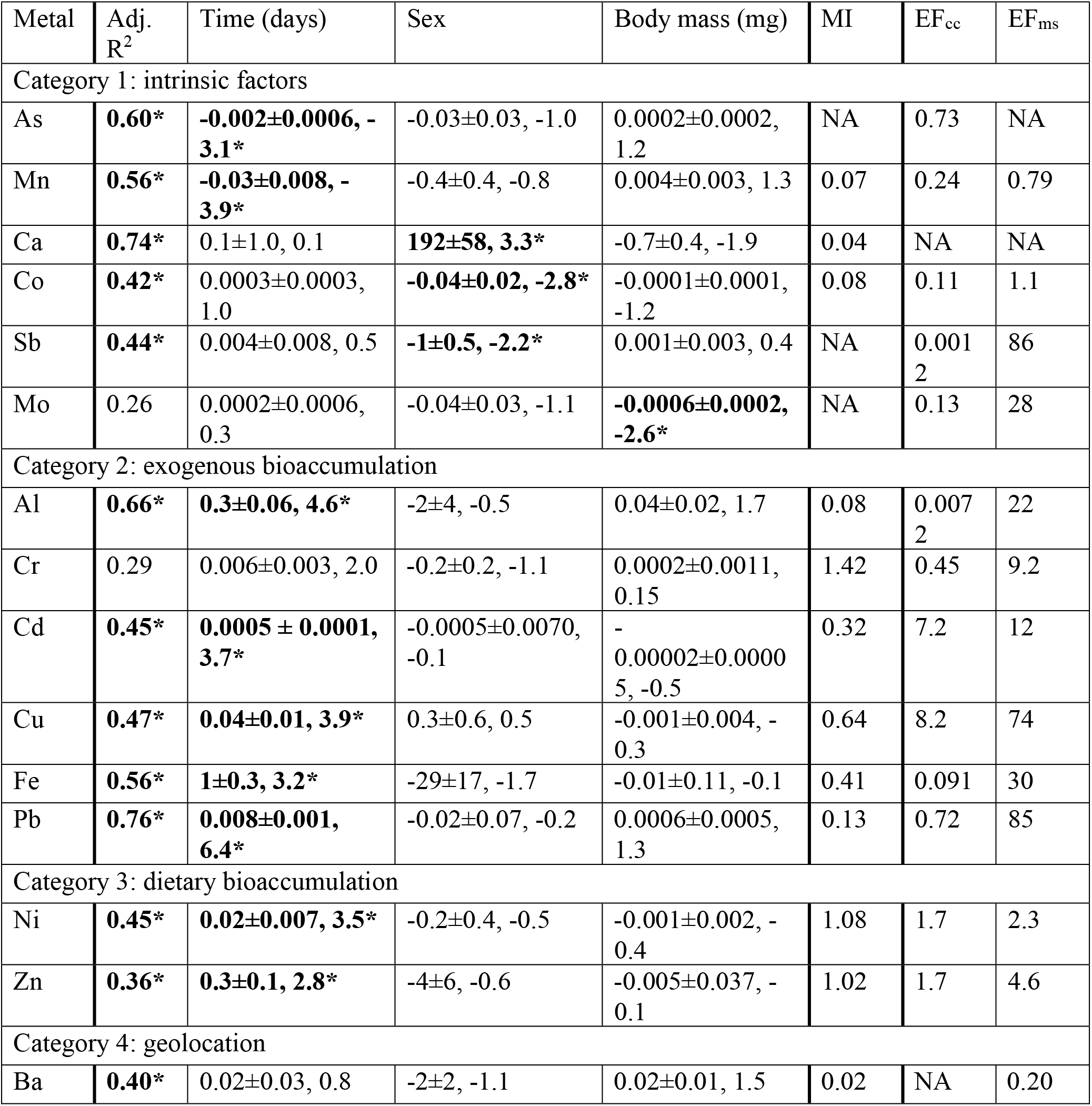
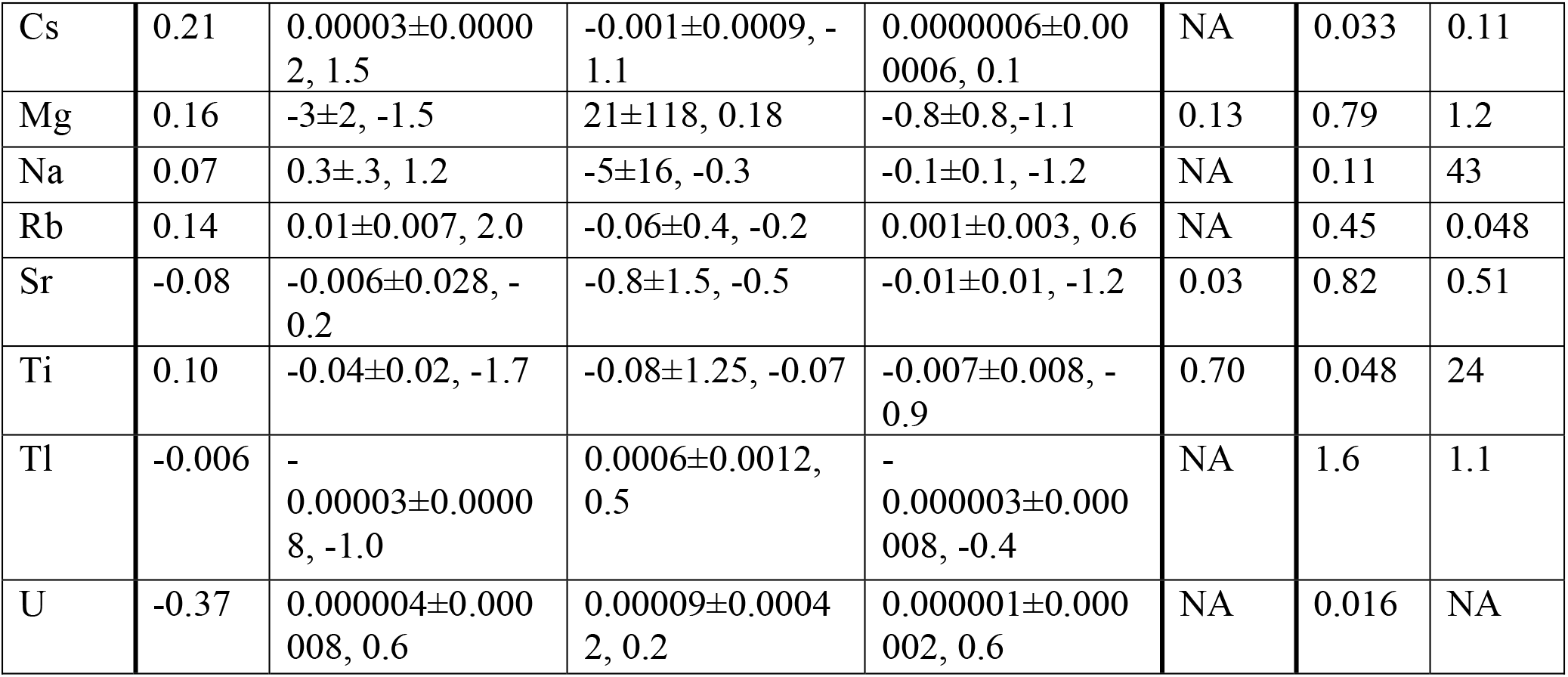
Metals are clustered into four categories based on their characteristics in the diet-switching experiment. Results of the multivariate multiple regression examining the contribution of time (days), sex, and body mass (mg) on metal concentrations of monarch wings in the diet-switching experiment are reported. Non-standardized (ng/mg) effect sizes are reported for each predictor (β ± SE, t-value); significant relationships are starred and in bold (α = 0.05). The sample size is 18 for all metals except As (n = 14), Tl (n = 16), and U (n = 10). The mobility index (MI), enrichment factor between the monarch wings and the continental crust (EF_cc_), and the enrichment factor between the monarch wings and the adult diet (EF_ms_) are also reported for each metal.

**Figure 2.**
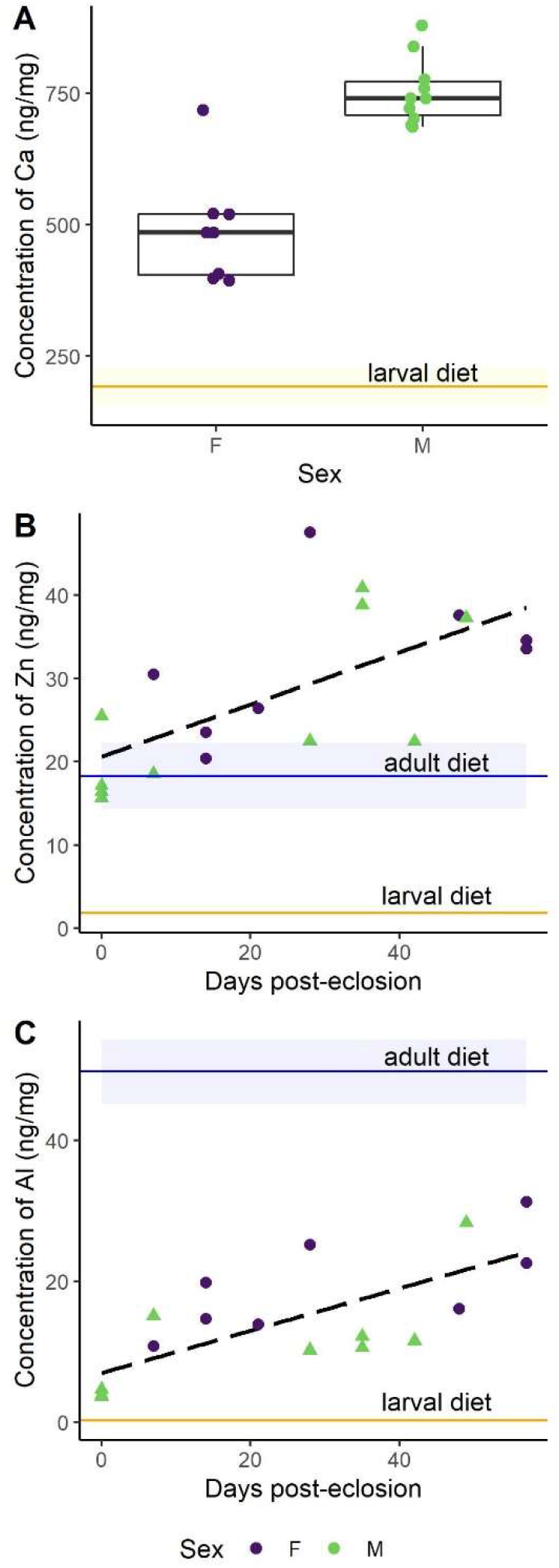
Metal concentration in monarch wings from the diet-switching experiment for representative metals classified in different functional categories. Horizontal blue lines depict the mean concentration (± SD) in the larval diet, milkweed (n = 3), and horizontal orange lines represent the mean concentration (± SD) in the adult diet, maple syrup (n = 4). **(A)** Category 1: Boxplot of Ca concentration (ng/mg) in monarch wings by sex (RSD = 4%). Sex-based differences were detected with males (green) having more Ca than females (purple). The milkweed concentration (18,500 ± 1800 ng/mg) is not shown as it far exceeds that of the monarch wings. **(B)** Category 2: The concentration of Zn (ng/mg) in monarch wings increased over the 8 weeks (RSD = 1%). **(C)** Category 3: The concentration of Al (ng/mg) in monarch wings increased over the 8 weeks (RSD = 2%).

Enrichment factors and mobility indices calculated for each of the 23 metals showed distinctive patterns in the concentrations between the monarch butterfly wings and the larval diet (i.e., milkweed), the adult diet (i.e., maple syrup), and a proxy for mineral dust (i.e., continental crust). Chromium was the only metal with a mobility index greater than one indicating that it is biomagnified in monarch wings compared to milkweed (Table 1). Nickel and Zn had mobility indices close to one. The remaining elements showed low concentrations in monarch wings compared to milkweed indicating biodilution of these metals. The lowest mobility indices were found among the alkaline earth metals (i.e., Mg (0.13), Ca (0.04), Sr (0.03), and Ba (0.02)). Enrichment factors less than one suggest that the metal was susceptible to contamination from either the adult diet (EF_ms_) or mineral dust (EF_cc_). Of the metals that showed an increase in concentration over time, Al and Fe showed particularly low EF_cc_, indicating that the increase in concentration could be due to additions of Al and Fe from mineral dust. Comparatively low EF_ms_ for Ni and Zn suggest that the bioaccumulation seen in these metals may be due to additions from the adult diet.

Based on the results of the PCAs, multiple multivariate regression, mobility indices, and enrichment factors, metals were separated into four categories: (1) metals affected by intrinsic factors (e.g., sex, body mass), (2) metals showing bioaccumulation likely from external sources, (3) metals showing bioaccumulation likely from dietary sources, and (4) metals without significant effects of intrinsic factors that maintain the initial metal concentration over time (Table 1). Metals were categorized as Category 1 if sex or mass were associated with metal concentration or there was a decrease in concentration over time. A few of these metals (i.e., Ca, Mn) were the main drivers of PC2 (Figures 1A and S1B). Metals assigned to Category 1 were As, Ca, Co, Mn, Mo, and Sb. For example, Ca was significantly more abundant in male wings than in female wings (Figure 2A), but no effect of time was detected. Category 2 metals had a positive association with time and relatively low EF_cc_, indicating that their increases in concentration over time were more likely to be sourced from mineral dust rather than from the adult diet. For instance, despite low Al levels in the maple syrup (EF_ms_ = 22), the concentration of Al in the monarch wings increased over time (Table 1). As a terrigenous metal, it is more likely that the accumulated Al is sourced from exogenous mineral dust (EC_cc_ = 0.0072). Some of the Category 2 elements are terrigenous metals, including Al and Fe, but others are associated with anthropogenic pollution (i.e., Cd, Cu, Cr, Pb). Several of the Category 2 metals are essential to living organisms (i.e., Cr, Cu, Fe), but others are toxic to animals even at low concentrations (i.e., Al, Cd, Pb). Category 3 metals (i.e., Ni, Zn) also had a positive association with time, but had high EF_cc_ and relatively low EF_ms_ and are therefore potentially contaminated endogenously through the adult diet. As an illustration, the monarch wings showed a two-fold accumulation of Zn over time from a concentration similar to that of the larval diet (Figure 1B). The level of this essential metal was lower in the adult diet than in the larval diet, but the EF_ms_ was relatively low (4.6), suggesting that the increase in Zn may be sourced, at least partially, from the adult diet. Finally, metals were assigned to Category 4 if they showed no significant associations with time, sex, or body mass. These metals were Ba, Cs, Mg, Na, Rb, Sr, Ti, Tl, and U.

### 3.2 Strontium and Strontium isotope ratios

The Sr and ^87^Sr/^86^Sr of the diet-switching experiment monarch wings, larval diet, and adult diet are reported in Table S2. Based on its characteristics, Sr was considered as Category 4 (Table 1), a metal that does not bioaccumulate and is not significantly affected by sex or body mass. The mobility index (0.03) indicated that Sr is biodiluted in the monarch wings. These low concentrations did not have significant associations with time, sex, or body mass (Table 1), although a non-significant sex-based pattern could be observed (Figure 3A).

**Figure 3.**
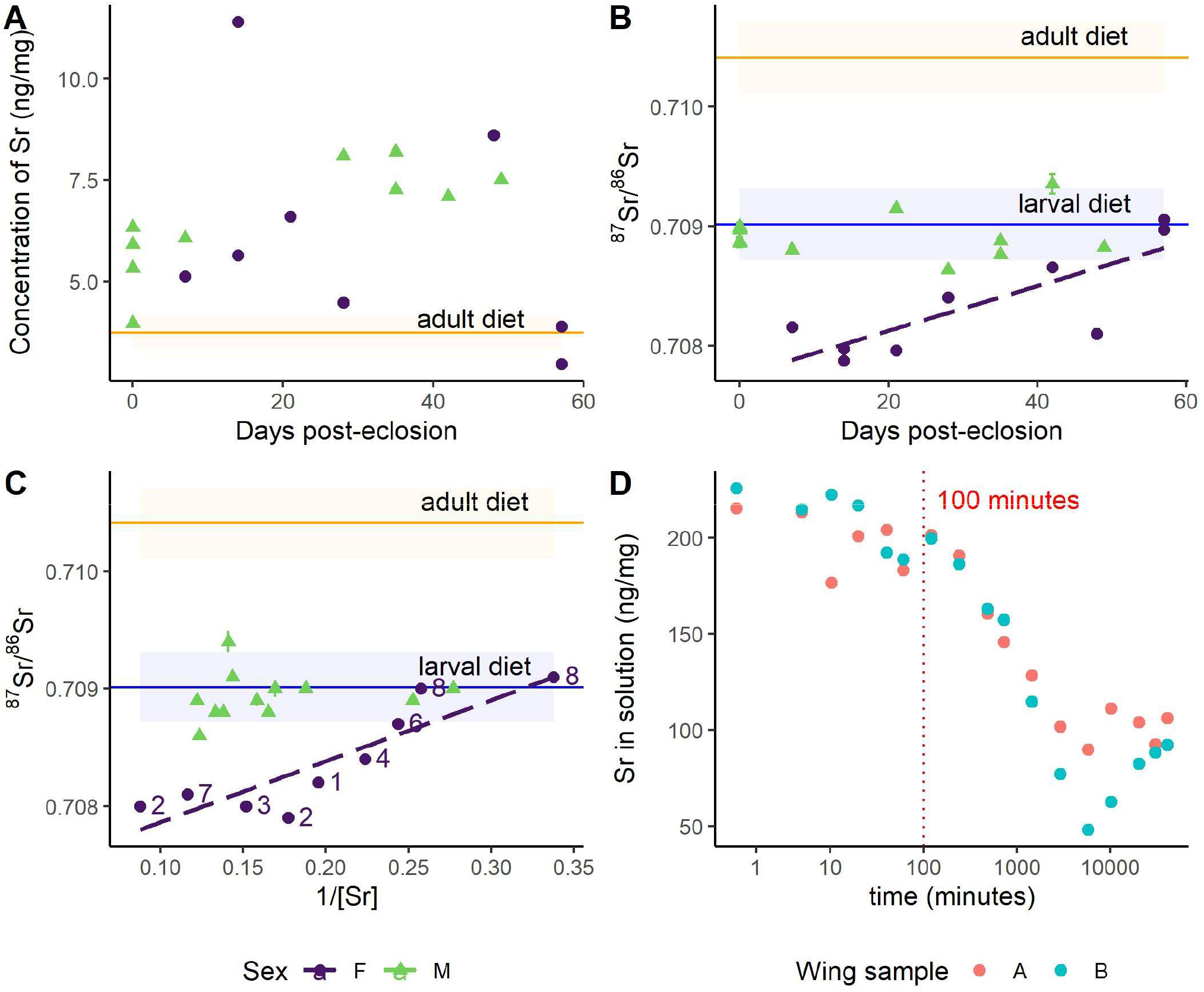
Strontium and strontium isotope ratios in monarch wings during the diet-switching and wetting experiments. Horizontal blue lines depict the mean concentration (± SD) in the larval diet, milkweed (n = 3), and horizontal orange lines represent the mean concentration (± SD) in the adult diet, maple syrup (n = 4). **(A)** Concentration of Sr (ng/mg) in monarch wings over 8 weeks (RSD = 1%). The concentration of Sr in the larval diet is not shown as it far exceeds that of the monarchs (189 ± 21 ng/mg). **(B)** Strontium isotope ratios (± SE) in monarch wings over 8 weeks; the ^87^Sr/^86^Sr ratios of the female monarch wings show a linear relationship with time. **(C)** Strontium isotope ratios (± SE) against the inverse of strontium concentration. The significant linear relationship for females is displayed; numbers depict the weeks post-eclosion that the female monarch was sampled. **(D)** Concentration of Sr (ng/mg) in solution through time during the wing wetting experiment (n = 2).

Multiple regression of monarch wing ^87^Sr/^86^Sr was performed with predictors time (days), sex, the inverse of Sr (mg/ng), and interactions between time and sex and the inverse of Sr and sex (adj. R^2^ = 0.79). As expected, the range of male ^87^Sr/^86^Sr matched well with the ^87^Sr/^86^Sr of the larval diet and did not change over time (p = 0.2; Figure 3B) or with the inverse of Sr (p =0.2; Figure 3C). However, on average, the wings of the female monarchs were 0.0014 less radiogenic than the males (β = 0.0014 ± 0.0004, 3.6, p < 0.01) and the ^87^Sr/^86^Sr of the female wings increased slowly over time (β = 0.000010 ± 0.000004, 2.3, p = 0.03), although the lack of female monarchs sampled at Week 0 may have been biasing this result (Figure 3B). If the adult diet is responsible for the increasing ^87^Sr/8^86^Sr of the female monarch wings over time, it seems unlikely that it is through accumulation, as no positive association between Sr and time was detected (p = 0.8; Table 1). The ^87^Sr/^86^Sr of the female monarch wings was associated with the inverse of Sr (β = 0.0037 ± 0.0011, 3.4, p < 0.01), such that females with higher concentrations of Sr had ^87^Sr/^86^Sr more disparate from those of the males and larval diet (Figure 3C). Given that radiogenic strontium isotopes do not fractionate but instead indicate sources of Sr, the principles of mixing models can be used to find missing endmembers (i.e., the ^87^Sr/^86^Sr of pure sources of Sr to the monarch wings). The intercept of regression indicated a missing endmember for the female monarchs with a less radiogenic ^87^Sr/^86^Sr of about 0.7073. Females with higher abundances of Sr in their wings tended to be those sampled earlier in the experiment; lower concentrations of Sr in the Week 8 female monarch wings aligned them with the expected ^87^Sr/^86^Sr of the larval diet (Figure 3C). Thus, the female monarchs were likely exposed to the less radiogenic Sr source either as larvae or as young adults (i.e., less than a week old).

The wetting experiment shows that exogenous contamination of Sr in butterfly wings by wetting is possible. After we submerged the monarch wing in the aqueous solution, we detected substantial changes in the Sr concentration of the solution after 100 min. (Figure 3D). A new equilibrium was reached after about a week. Since the amount of Sr in the solution decreased over time, it is likely that the Sr accumulated in the wing.

### 3.3 Lead and Lead isotope ratios

Lead was categorized into Category 2 because Pb concentrations in the monarch wings increased over time, likely sourced from mineral dust (Table 1; Figure 4A). Of all the toxic metals, only Pb levels in the milkweed (0.22 ± 0.06 ng/mg, n = 3) and monarch wings (0.30 ± 0.20 ng/mg, n = 18) were found to exceed the permissible limits of human food (i.e., 0.01-1 ng/mg; FAO and WHO 2019), but Pb levels in the maple syrup (1.1 ± 0.6 ng/g, n = 4) were well below health guidelines. Lead isotope ratios showed that the larval diet matched well with regional environmental ^208^Pb/^206^Pb and ^207^Pb/^206^Pb signatures measured in snowpack (Figure 4B; Simonetti *et al*. 2000a, 2000b). The early monarchs (i.e., Week 0) also matched with the larval and environmental Pb isotopes, but within a week, the Pb isotopes of the monarch wings deviated to a distinct signature with higher ^208^Pb/^206^Pb and ^207^Pb/^206^Pb. Although the Pb isotope ratios were measured a year after the Pb levels, repeated measures showed that storage did not significantly change the initial Pb abundances (paired t-test: t = −1.3, df = 16, p = 0.20).

**Figure 4.**
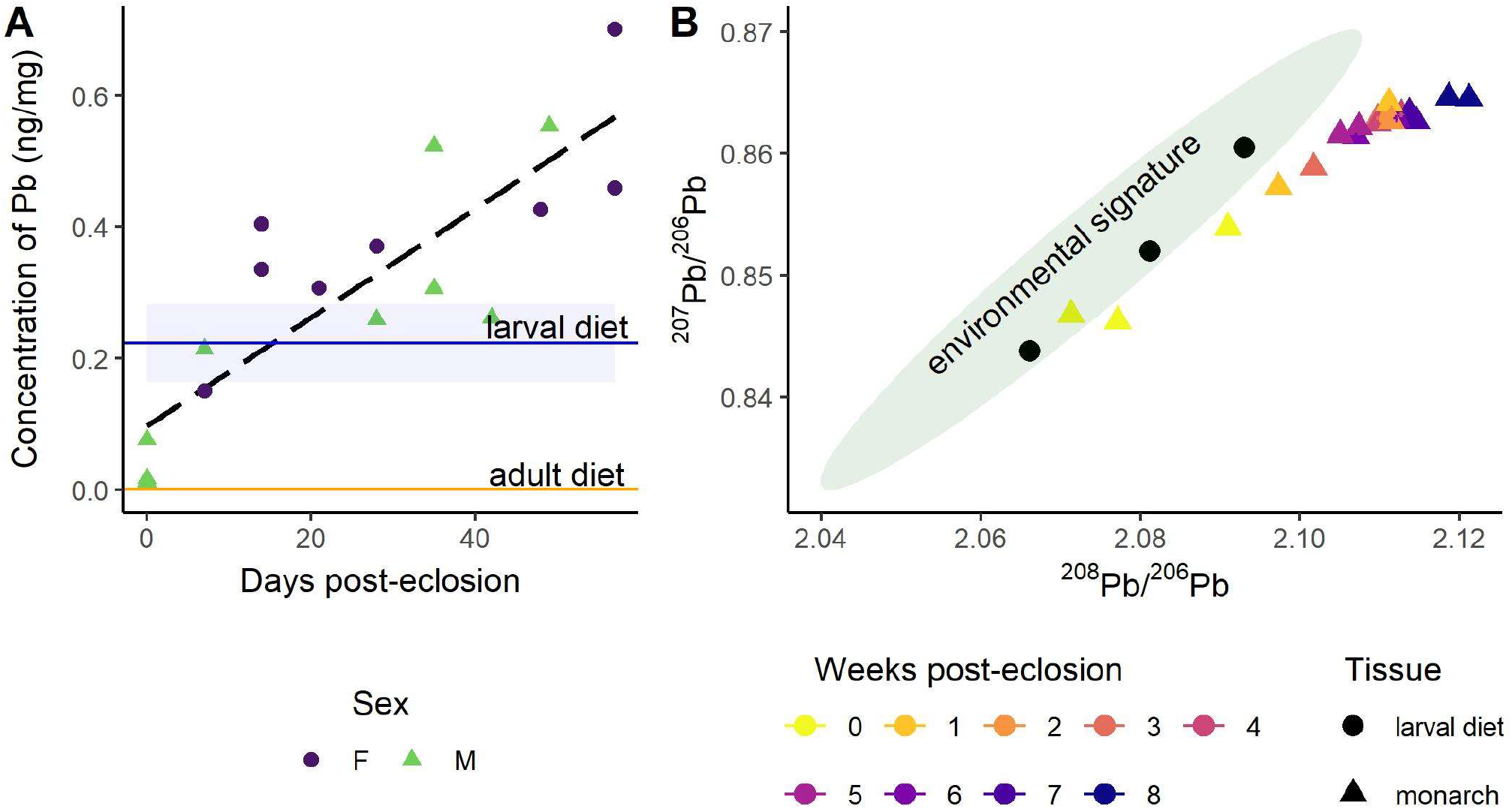
Change in lead and lead isotope ratios in monarch wings during the diet-switching experiment **(A)** Concentration of Pb (ng/mg) in monarch wings over 8 weeks (RSD = 1%); the linear relationship (dashed black line) with time is for all samples. The horizontal blue line depicts the mean concentration of Pb (± SD) in the larval diet, milkweed (n = 3), and the horizontal orange line represents the mean concentration of Pb (± SD) in the adult diet, maple syrup (n = 4). **(B)** Biplot of ^207^Pb/^206^Pb and ^208^Pb/^206^Pb ratios. Monarch wings are depicted as triangles with colors representing the age of the butterfly in weeks; the standard error is smaller than the symbols. Measurements of the larval diet are depicted as black dots (n = 3). The green 95% data ellipse shows the range of regional lead isotope ratios measured in snowpack (Simonetti *et al*. 2000b, 2000a).

### 3.4 Natural metal concentrations in lab-raised and wild monarchs

In this lab-based study, we controlled the environmental conditions of the monarch caterpillars and adults, minimized conspecific interactions, reduced flight capacity, and fed the adults a non-natural and choice-restricted diet. Thus, the variation in metal concentrations in the laboratory monarch wings is expected to be different from that of wild monarch wings. These differences can clarify important processes. Comparisons with wild monarch wings show that wing concentrations were similar for Cr, Mn, and Ni (Figure S5). Conversely, Mg, Al, Co, Cd, Ba, Pb had higher concentrations in the wild monarch wings and Cu, Zn, and Sr had lower concentrations in the wild monarchs (Figure S5).

## 4 Discussion

### 4.1 Regulation of metals in insect wings

The 23 metals analyzed in the diet-switching experiment were classified into four categories based on whether they were influenced by intrinsic factors like sex (Category 1), bioaccumulated metals from external sources (Category 2), bioaccumulated metals from dietary sources (Category 3), or maintained the natal signature (Category 4). Metals classified in Category 1, including As, Mn, Mo, and Ca, showed a decreasing concentration in the wings through time or a correlation with the intrinsic characteristics of individuals. These observations suggest that the levels of these metals are regulated by the insect. Category 1 metals were grouped into two sub-categories based on their distinct trends during the experiment (Table 1). The first sub-category included As and Mn, which showed decreasing concentrations in wings throughout the adult life (Table 1), suggesting that monarchs may reallocate these metals from the wings to other parts of the body or excrete them (Tibbett *et al*. 2021). Manganese is an essential co-factor in many enzymes but can impact insect behavior at low doses and, like all metals, is toxic in excess (Ben-Shahar 2018). Caterpillars of *Lymantria dispar* and *Cabera pusaria* have been shown to excrete excess Mn through frass and molting (Kula *et al*. 2014, Martinek *et al*. 2020), but adult mechanisms of excretion have not been shown and Mn homeostasis has not been studied in monarchs. Alternatively, As and Mn could be more concentrated in wing scales and decrease through time in the entire wing due to the progressive abrasion of scales as the individual ages.

The second sub-category of Category 1 included several metals (i.e., Ca, Co, Mo, Sb) that showed an influence of sex and/or body mass. However, sex and body mass were strongly correlated, with females being consistently heavier than males, making it challenging to disentangle the direct influence of mass from that of sex (Figure S3). Molybdenum showed a negative correlation to body mass (Table 1). Molybdenum is an essential metal that acts as a co-factor for many oxidase enzymes (Dow 2017), some of which relate to wing pigmentation (Feindt *et al*. 2018) and immune processes (Selvey 2020). The decrease in Mo concentration in wings with increasing body size might be related to differential regulation of the metal relative to body size, energy reserves, or sex (Orłowski *et al*. 2020). Alternatively, the decrease in Mo with increasing body size could be driven by differences in surface area-to-mass ratios. Female monarchs have been noted to have thicker wings than males (Davis and Holden 2015), leading to lower surface area-to-mass ratios in the wings. If Mo is predominantly found at the surface of the wing (e.g., pigmentation), this lower surface area-to-mass ratio could explain the pattern seen in this experiment (Kowalski *et al*. 1989). Similarly, we found that male monarch wings had significantly more Ca than females (Figure 2A). Notwithstanding the confounding effect of body mass, sex-based differences in Ca have also been detected in whole-body samples of butterflies (Dempster *et al*. 1986) and damselfly wings (Stuhr *et al*. 2018), suggesting that sex is the main driver of the observed differences in Ca in wings. Calcium is a known component of insect chorion (Studier 1996, Nickles *et al*. 2002) and reproductive organs (Clark 1958), and Ca has been found to be preferentially allocated to reproductive organs (Mesjasz-Przybyłowicz *et al*. 2014). This could explain the lower concentrations of Ca found in the wings of female monarchs, as it is possible that Ca is preferentially allocated or relocated from the wings to reproductive organs for oogenesis.

Little is known about the pathways of metal cycling and regulation in insects. Metallomics, the study of the metal-containing biomolecules that an organism uses, aims to identify and compare metal utilization, function, evolutionary trends, and interactions (Ogra and Hirata 2017). Of the six metals in Category 1 (Table 1), only the regulatory pathways of Ca, Co, and Mo have been well-characterized (Dow 2017, Zhang *et al*. 2019), illustrating the large knowledge gap that the newly-developing field of metallomics is seeking to fill. Metal isotopes that fractionate with physiological processes can be particularly useful to advance understanding but have been under-utilized in insects thus far (but see Nitzsche *et al*. 2020, 2022). For example, in vertebrates, calcium isotopes have been effectively used to investigate the modifications of Ca homeostasis due to pregnancy, lactation, and egg-laying (Tacail *et al*. 2020). Analyzing calcium isotope ratios of different insect tissues could shed light on Ca homeostasis in insects, such as the role of calcium-storing spherites (Dow 2017).

### 4.2 Bioaccumulation of metals in insect wings

We identified a suite of metals, including highly toxic (e.g., Al, Cd, Pb) and essential metals (e.g., Zn, Fe), that accumulated in the wings of the monarch butterflies over time (Categories 2 and 3; Table 1). Metals classified in Category 2 (i.e., Al, Cr, Cd, Cu, Fe, and Pb) increased in concentration over time in the monarch wings and had low enrichment factors between the monarch wings and continental crust (Table 1). These observations indicate that these metals were sourced, at least partially, from external sources such as mineral dust. For example, Fe and Al are highly concentrated in terrigenous dust, and previous studies on animal hair have shown that these elements were primarily sourced from exogenous sources in keratin tissues (Kempson *et al*. 2006, Hu *et al*. 2018). In keratin, dust and aerosol particles containing these metals accumulate on the surface of the tissue, and when wetted, these metals diffuse into keratin slowly. This pattern of diffusion is evidenced by the gradient of concentrations of these metals in keratinous tissue, from very high concentrations on the surface to low concentrations deeper in the tissue (Hu *et al*. 2018). In insects, high concentrations of Al in the cuticle of honeybees have also been observed, suggesting similar exogenous contamination (Shaw *et al*. 2018). However, given that butterfly wings are living tissue and that Fe is an essential element with known regulatory pathways (Slobodian *et al*. 2021), there are alternate explanations for the increase of Fe in wings through time. First, our experiment examined a single body part and therefore cannot comment on systemic metal homeostasis (Orłowski *et al*. 2020); Fe could be transported from different parts of the insect body for sequestration in response to perturbations to systemic homeostasis. Additionally, most of the accumulation of Fe in wings was observed between Week 0 and Week 1 (Figure 1B) and could be related to sclerotization and maturation processes as the wings finish forming within a few days post-eclosion (Honegger *et al*. 2008). Finally, Fe in wings could increase due to changes in Fe requirements over time, as has been observed in other insects (Shaw *et al*. 2018, Andreani *et al*. 2021). Metamorphosis is an energetically-costly process, and the teneral butterfly could be Fe-limited, causing a short-term rapid uptake of Fe to reach a regulatory threshold within the first week in an accumulator-regulator dynamic (Tibbett *et al*. 2021).

Further studies could use natural metal isotopes or isotope tracing to help constrain metal sources in insect tissues, as we exemplify in this study with the use of Pb isotopes (see discussion below). Similar to Al and Fe, Pb accumulated rapidly in the monarch wings by about 0.05 ng/mg per week. In parallel, we observed a temporal shift in Pb isotope ratios of the wings from a composition matching the local environment of the larval diet to a novel isotopic composition (Figure 4B). This indicated that the accumulating Pb is likely from external sources of atmospheric deposition and dust present in the laboratory air, such as from lead paint, pipes, or non-local mineral dust from a rock preparation laboratory located in the same building. Our work supports previous studies showing the contamination of insect tissues by exogenous Pb sources (e.g., Zhou *et al*. 2018, Smith and Weis 2020). The amount of Pb in the wings of these laboratory-reared monarchs exceeded permissible limits for human food, which is between 0.01 and 1 ng/mg (FAO and WHO 2019). Similarly, Pb in the wings of wild monarchs was also found to be above permissible limits (Table S4) to a greater extent than the laboratory monarchs (F(1, 116) = 24, p < 0.001); our experimental results suggest that the high Pb levels accumulated in wild monarch wings are likely sourced from anthropogenic pollution sources during their migration. Lead is toxic to insects and can cause mortality and sub-lethal effects such as reduced growth rate, body size, fecundity, and locomotor activity (Hirsch *et al*. 2003, Di *et al*. 2016, Zhou *et al*. 2021). Anthropogenic sources of Pb have unique Pb isotope ratios; thus, Pb and Pb isotope ratios of biological tissues could be used as indicators of environmental contamination sources (Smith *et al*. 2021). This principle has been applied to assess the local pollution exposure of non-dispersing insects, such as honeybees (e.g., Zhou *et al*. 2018, Smith and Weis 2020). We argue that Pb and Pb isotope ratios could also be used as bioindicators for migratory insects. As an individual insect moves along a migratory path, it will accumulate Pb from the different atmospheric pollution sources it encounters along the way. Measuring Pb isotope ratios in migratory butterflies could provide a tool to reconstruct migratory paths and assess the exposure to pollution of individuals going through this route. This tool would be key to relating migratory success and pollution exposure and could also validate migration routes reconstructed through pollen metabarcoding (Suchan *et al*. 2018) and wind trajectory modeling (Otuka *et al*. 2005). Ultimately, Pb isotopes in migratory insects could help identify key sources of pollution that are detrimental to specific species.

Metals classified in Category 3 (i.e., Ni and Zn) grouped metals which also bioaccumulated, but unlike Category 2, these metals were characterized by a relatively low enrichment factor between the monarch wings and maple syrup adult diet (Table 1). Nickel is not known to be an essential element to insects. Lepidopterans show variable effects to Ni exposure, from decreased body size and fecundity to no effect (Sun *et al*. 2013, Kobiela and Snell-rood 2018), suggesting that regulation mechanisms are species-specific (Tibbett *et al*. 2021). Nickel concentrations in the wings of monarchs in the diet-switching experiment did not significantly differ from those of wild monarchs (Figure S5; p = 0.4), suggesting that concentrations may be below regulatory thresholds and thus increasing due to non-specific uptake of metal from the adult diet. Conversely, Zn was found to be lower in wild monarch wings than in the diet-switching experiment (Figure S5; F(1, 116) = 40, p < 0.001), suggesting that the monarchs in the diet-switching experiment may have excess Zn. The high Zn in the laboratory monarch wings is unlikely to be purely a result of high Zn in the larval diet because the Zn levels in the milkweed provided in the diet-switching experiment (18 ± 4 ng/mg, n = 3; Table S2) are low compared to Zn concentrations measured in wild milkweed (10 - 66 ng/mg; Mitchell *et al*. 2020). Zinc is essential for insect reproduction and immunity (Cardoso-Jaime *et al*. 2022), and Zn homeostasis is known to be strictly controlled in insects (Xiao and Zhou 2016, Slobodian *et al*. 2021). However, excessive Zn can lead to decreased survival, growth rate, body size, and longevity (Jin *et al*. 2020). Although butterfly species are known to have different Zn tolerances, monarchs have been shown to be sensitive to Zn (Shephard *et al*. 2020). The accumulation of Zn seen in the diet-switching experiment could reflect an upregulation of Zn metalloproteins due to environmental or physiological conditions, such as increased metabolic processes (Butt *et al*. 2018). Zinc isotopes have been suggested as a tool to explore cycling and dietary sources of Zn to insects, but applications are currently hampered by the complicated pathways of Zn homeostasis and Zn fractionation in the body (Evans *et al*. 2017, Wanty *et al*. 2017, Nitzsche *et al*. 2020), which are poorly understood in insects compared to humans (e.g., Tanaka and Hirata 2018).

### 4.3 Geolocation using metals and metal isotopes

Although metals and metal isotopes that bioaccumulate have the potential to impart geographical information, as has been proposed for Pb isotopes, they are unsuitable for provenance studies that aim to estimate the natal origins of insects. Good candidate metals for the geolocation of migratory insects are those that (1) are unaltered in the tissues of dispersing insects, (2) have predictable spatial patterns, and (3) demonstrate a strong relationship between environmental and biological signatures. Nine metals belonging to Category 4 fit the first requirement and were unaltered over time in the monarch wings: Ba, Cs, Mg, Na, Rb, Sr, Ti, Tl, and U (Table 1). For example, the alkaline earth metals Sr and Mg showed consistent concentrations in the monarch wings over time, suggesting that the natal signature is conserved. Continuous-surface geolocation using metal concentrations would require spatial models of metal variation across the landscape (i.e., “metalscapes”). Mapping Sr and Mg of soil samples across the eastern USA demonstrated that these metals have distinctive spatial patterns, at least in the soil (Figure S6). However, correlations between metal concentrations in the soil and the biosphere (i.e., milkweed) were weak and non-significant (Table S5). Previous studies have also found similarly weak correlations in both natural (Wieringa *et al*. 2020) and controlled laboratory conditions (Lin *et al*. 2021). Exploration of possible predictors of Sr and Mg concentrations in milkweed highlighted land use, milkweed species, month, and year, but not soil concentration (Table S5). It is possible that some metal concentrations may vary at too fine a scale to be suitable for geolocation using continuous surfaces (Norris *et al*. 2007). Before metals can be used as geolocation biomarkers, further studies will need to improve the predictive power of spatial reference models.

Despite the apparent stability of the concentration of these metals from Category 4 through time, caution should be taken in applying these metals for insect geolocation. Of these metals, Mg and Na are essential metals and are thus expected to be regulated. Our results cannot differentiate between metals whose concentrations were unaltered in the monarch wings due to a lack of bioaccumulation and those that were held at a stable concentration through regulation for metal homeostasis. In the case of homeostasis, the metal concentration may be altered under different stressors not experienced in this experiment, such as starvation, disease, different environmental conditions, or increases in metal exposure. These stressors may alter the ability of the organism to maintain homeostasis (Hopkin 1990) and thus disrupt the apparent larval signature. Similarly, not all sources of potential exogenous contamination were explored in this study. For example, we showed that Sr in monarch wings is susceptible to contamination or exchange after about 100 min. when submerged in Sr-concentrated aqueous solutions (Figure 3D). This supports the finding that Sr is soluble and that exogenous contamination of Sr in keratinous hair readily occurs when hair is submerged in water (Hu *et al*. 2018, 2020). Monarchs are behaviorally averse to submergence in water and are unlikely to be exposed to these artificial conditions in nature (i.e., full submergence in highly Sr concentrated waters). However, they are likely to be exposed to rain which usually has a low Sr concentration, except near coastal areas, but can have isotopically variable signatures (Nakano *et al*. 2001). Further studies will need to investigate the effect on Sr concentration and ^87^Sr/^86^Sr of wetting from rainwater with varied concentrations and isotopic signatures. As evidenced by our wetting experiment, it would take over an hour of full submersion in Sr-concentrated water to exchange Sr in the wing because the superhydrophobic property on the surface of the wings limit the exchange of metals. However, the regular or sudden wetting by rain during storms could potentially alter, accumulate, or exchange some Sr in the wings, particularly in coastal regions where Sr is more concentrated in rainwater. Consequently, future studies should investigate the susceptibility of Sr and other metals to contamination under real-world conditions, including rain, aerosols, and pollution. Raising butterflies in the wild at specific sites with known metal and metal isotope composition could help reinforce the applicability of these tools for geolocation.

We have defined a suite of metals suitable for the geolocation of monarch butterflies, but it may be inappropriate to extrapolate these findings to other organisms, including other Lepidoptera. The specialized structures that aid in metal transport can differ between insect taxa, leading to phylogenetic-based differences in metal uptake, transfer, speciation, and segregation (Tibbett *et al*. 2021). This may be especially important for non-nectarivorous species because nectar contains low concentrations of metals compared to the diets of herbivores or carnivores. Caution should also be taken when using whole-body samples rather than a specific insect tissue with relatively low metabolic activity (Orłowski *et al*. 2020); it can be assumed that whole-body specimens will show greater changes over time than the wings due to greater inputs from the adult diet (Dempster *et al*. 1986). Ultimately, many sources of variability in metal composition need to be further explored, including the effects of host plant (Bowden *et al*. 1984), microbial communities (Consuegra *et al*. 2020), parasitism (Sures 2004), type of metamorphosis and life-stage, reproductive status, homeostatic mechanisms, and their interactions (Tibbett *et al*. 2021).

Strontium isotope ratios have already been applied to the geographic assignment of monarch butterflies and other lepidopteran species and showed great promise for increasing the precision of isotope-based geographic assignment (Reich *et al*. 2021, Dargent *et al*. 2022). Here, we confirmed the assumption that the ^87^Sr/^86^Sr of monarch wings remain the same over time and continue to represent the larval signature despite adult feeding, at least in males (Figure 3B). While we showed that the Sr in wings was susceptible to exchange in extreme wetting conditions (Figure 3D), we argue that these conditions are unlikely in nature and do not question the applicability of this geolocation tracer. More surprisingly, we also observed sex-based differences in ^87^Sr/^86^Sr, with females having a lower ^87^Sr/^86^Sr than both the males and their larval diet (Figures 3B and 3C). As ^87^Sr/^86^Sr trace the mixing of isotopically distinct sources (and not isotopic fractionation), this observation suggests that there is an unidentified source of Sr to the female monarchs. Strontium is chemically similar to Ca and is inadvertently substituted for calcium ions in the body (Capo *et al*. 1998). Based on our results that Ca and, to a lesser degree, Sr in wings vary by sex (Figures 2A and 3A), we argue that the difference in ^87^Sr/^86^Sr is potentially related to the preferential allocation of Ca (and thus Sr) to female reproductive organs for oogenesis, although no eggs were laid during the experiment. In this case, the ^87^Sr/^86^Sr of the wings would be determined by the milkweed ^87^Sr/^86^Sr from a discrete timestep rather than an integration of the entire larval stage. However, previous studies investigating ^87^Sr/^86^Sr in monarch wings did not detect any differences between males and females (Flockhart *et al*. 2015), and it is possible that our result reflects the low number of samples. Alternatively, the differences in ^87^Sr/^86^Sr between the sexes could be due to the high sensitivity of butterfly wings immediately after eclosion. The female monarchs emerged from their chrysalises one day earlier than the males (Figure S4). The day that the females eclosed coincided with an observation that paint odors could be smelled in the laboratory where the adult monarchs were kept, although the laboratory itself was not painted. Many indoor paints contain high concentrations of Sr (Van Gorkum and Bouwman 2005), as do gypsum-based mixtures (Huang 2020), which are often applied to fill holes and sanded smooth prior to painting. High concentrations of dust from either of these substances present on the day the female monarchs eclosed from their chrysalises could have deposited on the surface of the wings, and since butterfly wings are soft and wet on the day of eclosion, it is possible that external Sr could have mobilized and penetrated the wing tissues. We suggest that future studies explore the link between sex and ^87^Sr/^86^Sr in monarch wings, and the sensitivity of metals in butterfly wings to environmental conditions on the day of eclosion.

### 4.4 Conclusion

In summary, this controlled diet-switching experiment of monarch butterflies demonstrated the complexity and promises of using metals and metal isotopes in ecological studies. We showed that several metals and metal isotopes have clear potential to advance our understanding of insect ecology. First, six metals were regulated and/or changed with sex and body mass, reflecting their involvement in physiological processes (e.g., Mn, Mo, Ca). Other metals bioaccumulated in the insect wing either from the diet (i.e., Ni, Zn) or from external sources like mineral dust (e.g., Al, Fe, Pb). Finally, other metals remained stable through the life of the monarch (e.g., Sr, Mg) and are good potential candidates for developing geolocation tools. This study also demonstrated that ^87^Sr/^86^Sr ratios are a promising geolocation tool but that the potential for contamination of the natal signature during wetting events should be further explored. We also show that lead isotopes have a potential to trace the integrated pollution sources of migratory insects. Combining Pb isotopes and toxic heavy metal concentrations in insect tissues (e.g., Hg, Cd, As, Ni, Cu) is a promising avenue to better understand the mechanisms and sources of metal toxicity in insects. However, this study also demonstrated the complexity of metal cycling in insect tissues and the need for further work to fill this knowledge gap. Our results underline the need to understand metal cycling prior to using metals as geolocation tools in ecological studies. We argue that only a few metals have a potential for geolocation and that metals with changing concentrations through time should either be excluded or their incorporation mechanisms understood before using them as geolocation biomarkers. Even in cases where the adult insect does not feed and, thus, no dietary input is expected (e.g., eastern spruce budworm), metals which may be altered exogenously should be carefully evaluated.

## 5 Conflict of Interest

The authors declare that the research was conducted in the absence of any commercial or financial relationships that could be construed as a potential conflict of interest.

## 6 Author contributions

MSR conceived the idea for the study; MSR, FD, and HK designed the insect rearing component; MSR, CPB, MKK, FD, and LH designed the metal and metal isotope analysis component; DTTF and DRN collected the wild monarch and milkweed samples; MSR, MKK, LH, and CPB analyzed the data, MSR led the writing of the manuscript. All authors contributed to the drafts and gave final approval for publication.

## 7 Funding

This study was funded by the New Frontiers in Research Fund (CPB, GT) and Syngenta Canada, Inc. (DRN, MSR). GT acknowledges funding from the grant PID2020-117739GA-I00 from MCIN / AEI / 10.13039/501100011033. MSR was supported by the Queen Elizabeth II Graduate Scholarship in Science and Technology (QEII-GSST) and Ontario Graduate Scholarship.

## 8 Acknowledgements

Monarchs for the diet-switching experiment were collected under Ontario Ministry of Natural Resources collection authorization #1093661. We would like to thank Shuanquan Zhang for his expert guidance in operating the MC-ICP-MS at Carleton University, Joseph Spencer for running the Pb columns, and Emma Brown, Daniela Quintero, and Aldo Camilo Martinez-Becerril for assisting with animal husbandry.

## 9 Data Availability Statement

The original contributions presented in the study and the R code used in the analysis are archived with the Open Science Foundation (https://osf.io/n4pvm/?view_only=34c30a945e364029a655a9a4665f7a01).

## Notes

### Competing Interest Statement

The authors have declared no competing interest.

