## Supplementary Material for "Metals and metal isotopes in insect wings: Implications for diet, geolocation and pollution exposure"

### 1 Peach Leaves Standard

| **Table S1.** The mean, standard deviation, and relative standard deviation (calculated as sd/mean*100) as calculated for SRM 1547 Peach Leaves (National Institute of Standards and Technology) subjected to the same preparation procedure as the milkweed samples (i.e., ashed, digested). The samples (n = 3) were analyzed during three ICP-MS runs. Sample concentrations cannot be directly compared to reference values because the sample concentrations are calculated based on the mass of the sample after it is dry ashed, a process which removes carbon, hydrogen, nitrogen, and some oxygen and volatile elements (e.g., Cl, Br, S) from the sample. | | | | |
| --- | --- | --- | --- | --- |
| Element | mean (ng/mg) | SD | n | RSD (%) |
| Mg | 46156 | 5560 | 3 | 12 |
| Al | 2343 | 86 | 3 | 4 |
| Cr | 10 | 3 | 3 | 32 |
| Mn | 993 | 78 | 3 | 8 |
| Co | 0.84 | 0.13 | 3 | 15 |
| Ni | 12 | 7 | 3 | 56 |
| Cu | 53 | 23 | 3 | 44 |
| Zn | 236 | 61 | 3 | 26 |
| Sr | 608 | 158 | 3 | 26 |
| Cd | 0.24 | 0.05 | 3 | 21 |
| Pb | 7.6 | 0.4 | 3 | 6 |
| Ba | NA | NA | 0 | NA |

### 3 Diet-switching experiment

| 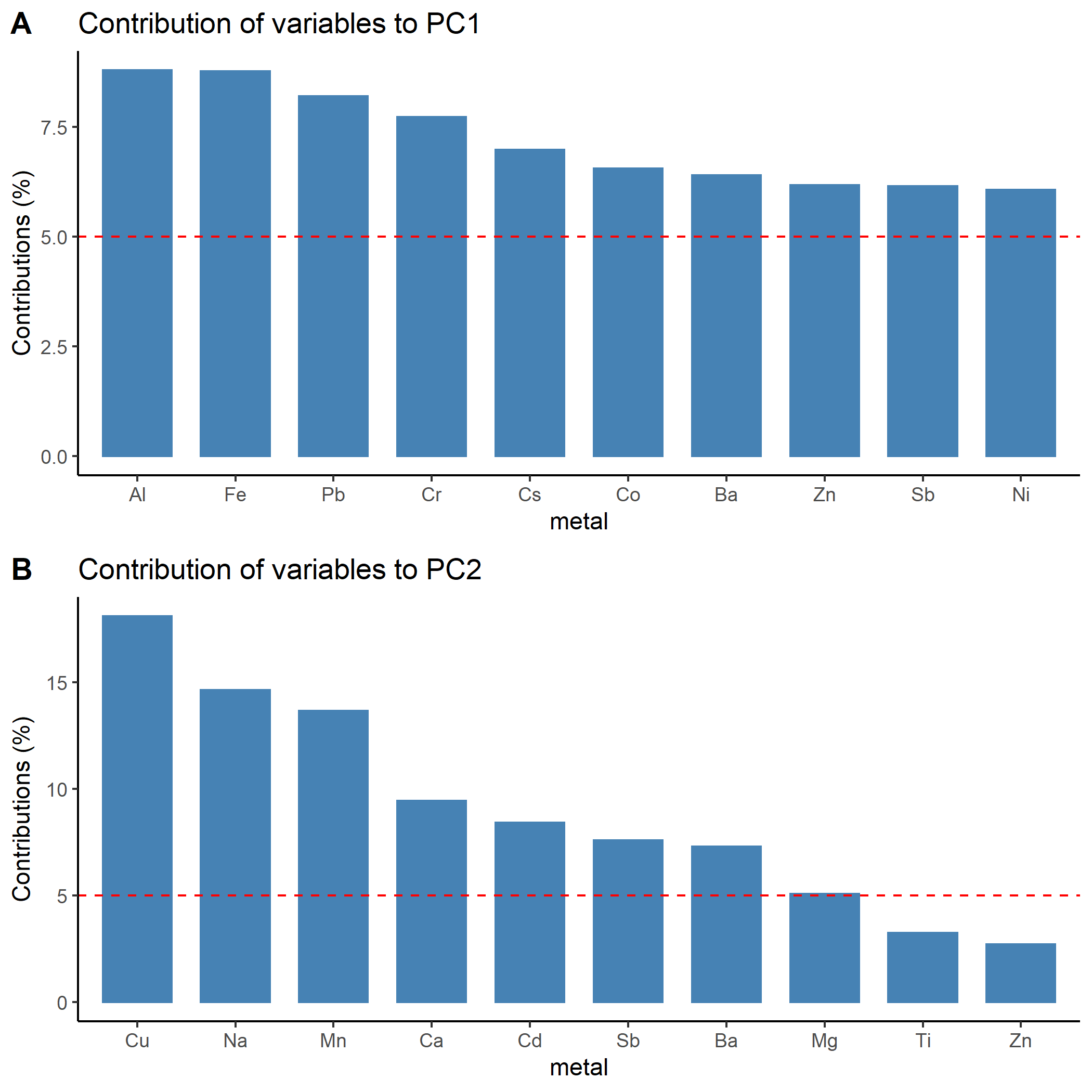 | **Figure S1** Contribution of metals in the wings of monarch butterflies from the diet-switching experiment to **(A)** PC1 and **(B)** PC2. Produced using the *factoextract* package in R (Kassambara and Mundt 2020) |
| --- | --- |

| **Table S2.** Metal concentrations (ng of metal per mg of sample material (mean ± SD (n)) of the larval diet, adult diet, and monarch butterfly wings at different weeks of the diet-switching experiment. | | | | | | | | | | | |
| --- | --- | --- | --- | --- | --- | --- | --- | --- | --- | --- | --- |
| metal | larval diet (n=3) | Monarch wings separated by week of sampling | | | | | | | | | adult diet (n=4) |
|  |  | 0 | 1 | 2 | 3 | 4 | 5 | 6 | 7 | 8 |  |
| Al | 50±5 | 3.9±0.5 (4) | 13±3 (2) | 17±4 (2) | 14 (1) | 17±11 (2) | 11±1 (2) | 12 (1) | 22±9 (2) | 27±6 (2) | 0.2±0.03 |
| As | NA | bdl | 0.17±0.08 (2) | 0.11±0.03 (2) | 0.13 (1) | 0.10±0.10 (2) | 0.04±0.01 (2) | 0.05 (1) | 0.04±0.01 (2) | 0.06±0.03 (2) | NA |
| Ba | 25±5 | 0.6±0.1 (4) | 5.0±2.5 (2) | 4.7±0.8 (2) | 5.1 (1) | 7.8±6.2 (2) | 3.0±1.3 (2) | 2.1 (1) | 3.3±0.5 (2) | 4.4±2.2 (2) | 5.4±0.6 |
| Ca | 18477±1802 | 718±26 (4) | 546±198 (2) | 558±226 (2) | 521 (1) | 622±194 (2) | 807±44 (2) | 880 (1) | 603±168 (2) | 457±89 (2) | 191±37 |
| Cd | 0.01±0.002 | 0.003±0.001 (4) | 0.011±0.002 (2) | 0.016±0.002 (2) | 0.008 (1) | 0.012±0.008 (2) | 0.025±0.002 (2) | 0.011 (1) | 0.036±0.027 (2) | 0.028±0.006 (2) | 0.0004±0.0001 |
| Co | 0.14±0.02 | 0.011±0.005 (4) | 0.042±0.007 (2) | 0.081±0.024 (2) | 0.073 (1) | 0.052±0.011 (2) | 0.046±0.005 (2) | 0.038 (1) | 0.085±0.030 (2) | 0.041±0.008 (2) | 0.013±0.001 |
| Cr | 0.5±0.2 | 0.7±0.1 (4) | 1.2±0.4 (2) | 0.9±0.2 (2) | 0.9 (1) | 1.2±0.3 (2) | 1.0±0.2 (2) | 1.1 (1) | 1.2±0.1 (2) | 1.1±0.05 (2) | 0.033±0.002 |
| Cs | NA | 0.0018±0.0008 (4) | 0.0035±0.0009 (2) | 0.0053±0.0003 (2) | 0.0045 (1) | 0.006±0.0002 (2) | 0.0041±0.0004 (2) | 0.005 (1) | 0.0041±0.0003 (2) | 0.0039±0.0005 (2) | 0.011±0.001 |
| Cu | 7.7±0.3 | 4.9±0.8 (4) | 4.2±0.1 (2) | 5.4±0.3 (2) | 5.1 (1) | 5.1±0.3 (2) | 7.2±0.3 (2) | 5.4 (1) | 7.0±0.5 (2) | 6.5±0.8 (2) | 0.02±0.03 |
| Fe | 112±6 | 45±8 (4) | 79±8 (2) | 97±9 (2) | 93 (1) | 92±59 (2) | 83±25 (2) | 84 (1) | 147±2 (2) | 108±17 (2) | NA |
| Mg | 4168±330 | 529±261 (4) | 160±9 (2) | 190±34 (2) | 193 (1) | 199±1 (2) | 260±2 (2) | 263 (1) | 292±56 (2) | 205±28 (2) | 73±5 |
| Mn | 79±20 | 5.1±0.6 (4) | 5.4±0.4 (2) | 4.5±0.4 (2) | 5.5 (1) | 4.4±1.2 (2) | 3.5±0.4 (2) | 3.0 (1) | 4.3±0.6 (2) | 3.9±0.7 (2) | 1.7±0.2 |
| Mo | NA | 0.36±0.06 (4) | 0.34±0.07 (2) | 0.32±0 (2) | 0.25 (1) | 0.37±0.05 (2) | 0.37±0.05 (2) | 0.4 (1) | 0.38±0.06 (2) | 0.34±0.08 (2) | 0.0038±0.0003 |
| Na | NA | 60±16 (4) | 46±24 (2) | 60±10 (2) | 60 (1) | 60±4 (2) | 106±42 (2) | 43 (1) | 79±17 (2) | 69±18 (2) | 0.46±0.03 |
| Ni | 1.11±0.06 | 1.2±0.5 (4) | 1.6±0.1 (2) | 1.8±0.7 (2) | 1.9 (1) | 2.3±0.1 (2) | 1.5±0.2 (2) | 2.6 (1) | 3.0±0.7 (2) | 2.4±0.1 (2) | 0.25±0.02 |
| Pb | 0.22±0.06 | 0.03±0.03 (4) | 0.18±0.05 (2) | 0.37±0.05 (2) | 0.31 (1) | 0.32±0.08 (2) | 0.41±0.15 (2) | 0.26 (1) | 0.49±0.09 (2) | 0.58±0.17 (2) | 0.0011±0.0006 |
| Rb | NA | 0.2±0.1 (4) | 0.8±0.3 (2) | 1.1±0.1 (2) | 1 (1) | 2.1±0.4 (2) | 0.8±0.3 (2) | 1.2 (1) | 0.9±0.2 (2) | 1.1±0.1 (2) | 5.8±0.3 |
| Sb | NA | 0.06±0.02 (4) | 1.48±0.17 (2) | 2.15±0.44 (2) | 1.65 (1) | 2.19±1.11 (2) | 1.07±0.02 (2) | 0.49 (1) | 1.06±0.69 (2) | 1.4±0.07 (2) | 0.004±0.001 |
| Sr | 189±21 | 5.4±1.0 (4) | 5.6±0.7 (2) | 8.5±4.1 (2) | 6.6 (1) | 6.3±2.6 (2) | 7.7±0.7 (2) | 7.1 (1) | 8.1±0.8 (2) | 3.4±0.7 (2) | 3.7±0.4 |
| Ti | 10.2±0.3 | 7.1±2.1 (4) | 3.0±0.02 (2) | 4.2±1.2 (2) | 2.9 (1) | 3.4±0.6 (2) | 3.8±0.2 (2) | 3.1 (1) | 4.4±0.3 (2) | 4.2±0.6 (2) | 0.06±0.01 |
| Tl | NA | 0.0039±0.0026 (3) | 0.0045±0.0031 (2) | 0.0033±0.0004 (2) | 0.0042 (1) | 0.0025±0.0012 (2) | 0.0051±0.0012 (2) | bdl | 0.0023±0.0002 (2) | 0.0023±0.0004 (2) | 0.001±0.0001 |
| U | NA | bdl | 0.0008±0.0001 (2) | 0.0011±0.0009 (2) | 0.0010 (1) | 0.0012 (1) | bdl | bdl | 0.0008±0.0003 (2) | 0.0013±0.0003 (2) | NA |
| Zn | 18±4 | 19±5 (4) | 25±8 (2) | 22±2 (2) | 26 (1) | 35±18 (2) | 40±1 (2) | 22 (1) | 37.5±0.2 (2) | 34.1±0.7 (2) | 1.84±0.07 |

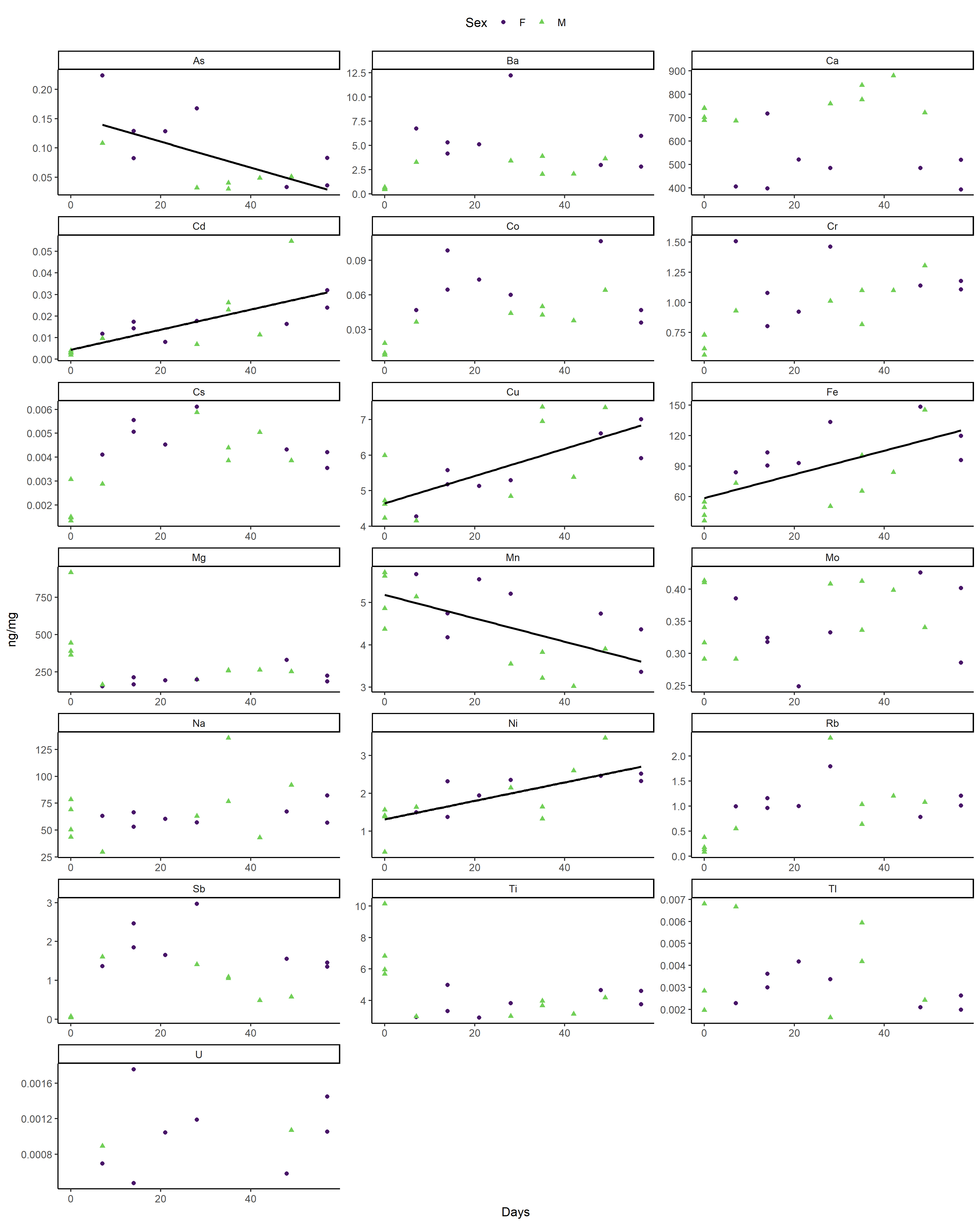

**Figure S2.** Metal concentrations (ng/mg) in monarch wings over 8 weeks from the diet-switching experiment. Males are indicated by green triangles and females by purple dots. Missing metals can be found in the main text (Zn, Al: Figure 2; Sr: Figure 3; Pb: Figure 4).

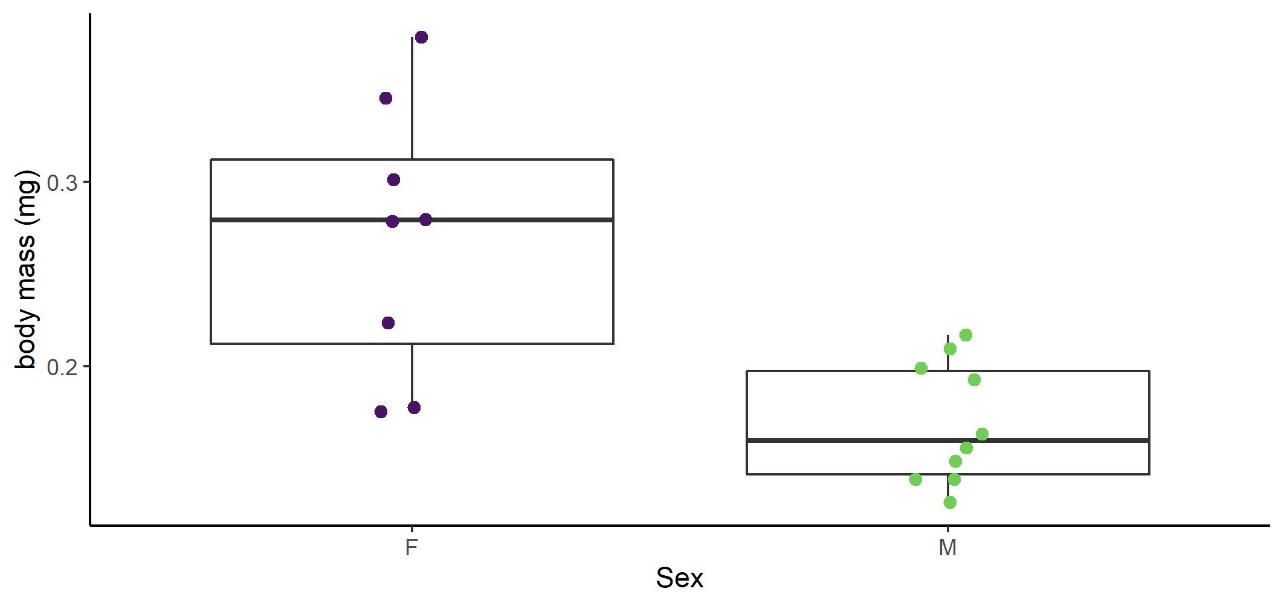

**Figure S3.** Boxplot demonstrating the confounding relationship between monarch body mass (mg) and sex (point-biserial correlation = -0 .70) for the diet-switching experiment.

| **Table S3.** A repeat of the multivariate multiple regression (Table 1) was performed for each metal in the wings of the monarchs from the diet-switching experiment. To test for model sensitivity to the confounding relationship between body mass and sex (Figure S3), sex was excluded from the regression, leaving time (days) and body mass (mg) as explanatory variables. Model fit is given as adjusted R^2^ and non-standardized (ng/mg) effect sizes are reported for each predictor (β ± SE, t-value); significant relationships are starred and in bold (α = 0.05). The sample size is 18 for all metals except As (n = 14), Tl (n = 16), and U (n = 10). Only a few regression results were sensitive to the removal of the sex variable: (1) for Ca and Sb, significant effects that were previously attributed to sex were now attributed to body mass, (2) the effect of body mass on Ba and Mn changed to statistically significant, and (3) the effect of days on Cr, Cs, and Rb became statistically significant. | | | |
| --- | --- | --- | --- |
| Metal | Adj. R^2^ | Time (days) | Body mass (mg) |
| Category 1: intrinsic factors | | | |
| As | **0.60*** | **-0.0018±0.0006, -2.9*** | 0.0003±0.0001, 2.8 |
| Ca | **0.57*** | -1.3±1.2, -1.1 | **-1.6±0.34, -4.8** |
| Co | 0.15 | 0.00061±0.00031, 2.0 | -0.00009±0.00008, -1.1 |
| Mn | **0.57*** | **-0.030±0.007, -3.9*** | **0.006±0.002, 2.9*** |
| Mo | **0.25*** | 0.0004±0.0006, 0.7 | **-0.0004±0.0002, -2.7*** |
| Sb | **0.29*** | 0.011±0.008, 1.4 | **0.006±0.002, 2.7*** |
| Category 2: exogenous bioaccumulation | | | |
| Al | **0.68*** | **0.30±0.06, 5.4*** | 0.05±0.02, 3.1 |
| Cr | **0.28*** | **0.007±0.003, 2.6*** | 0.001±0.0007, 1.4 |
| Cd | **0.49*** | **0.0005 ± 0.0001, 4.2*** | -0.00002±0.00003, -0.7 |
| Cu | **0.50*** | **0.038±0.009, 4.2*** | -0.002±0.003, -1.0 |
| Fe | **0.50*** | **1.2±0.3, 4.1*** | -0.13±0.08, -1.6 |
| Pb | **0.77*** | **0.008±0.001, 7.4*** | 0.0007±0.0003, 2.3 |
| Category 3: dietary bioaccumulation | | | |
| Ni | **0.47*** | **0.024±0.006, 4.2*** | -0.0002±0.0016, -0.1 |
| Zn | **0.38*** | **0.31±0.09, 3.5*** | 0.01±0.02, 0.5 |
| Category 4: geolocation | | | |
| Ba | **0.39** | 0.04±0.03, 1.4 | **0.024±0.007, 3.4*** |
| Cs | 0.20 | **0.00003±0.00002, 2.2*** | 0.000006±0.000004, 1.3 |
| Mg | 0.21 | -3.4±1.9, -1.8 | -0.95±0.52,-1.8 |
| Na | 0.13 | 0.37±.26, 1.4 | -0.11±0.07, -1.5 |
| Rb | 0.20 | **0.014±0.006, 2.3*** | 0.002±0.002, 1.0 |
| Sr | -0.02 | 0.0005±0.0245, -0.02 | -0.008±0.007, -1.2 |
| Ti | 0.16 | -0.04±0.02, -1.9 | -0.007±0.005, -1.3 |
| Tl | 0.05 | -0.00003±0.00002, -1.4 | -0.000006±0.000006, -1.0 |
| U | -0.18 | 0.000004±0.000007, 0.6 | 0.000001±0.000002, 0.6 |

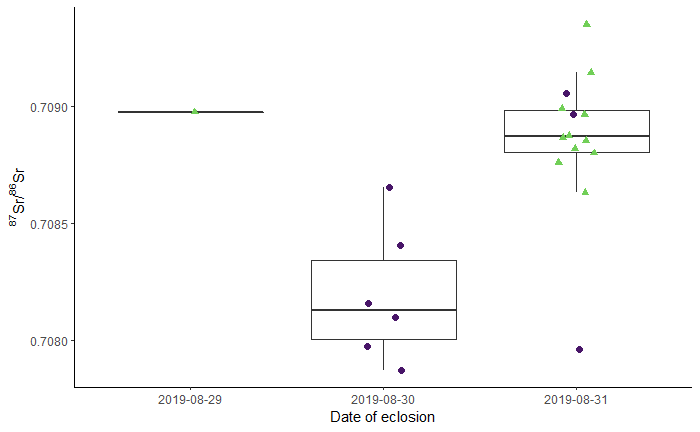

**Figure S4** Boxplot of ⁸⁷Sr/⁸⁶Sr by date of eclosion for the diet-switching experiment. Most male monarchs (green triangles) emerged from their chrysalises one day after the females (purple dots).

### 4 Natural metal concentrations in ashed milkweed and monarch wings

#### 4.1 Elemental analysis of ashed milkweed samples

To estimate the elemental concentrations of wild milkweed plants of different species (*Asclepias* spp.) in eastern North America, 85 milkweed samples were analyzed for a suite of 12 elements (i.e., Al, Ba, Cd, Co, Cr, Cu, Mg, Mn, Ni, Pb, Sr, and Zn). The milkweed samples were collected and analyzed as part of a recent study (Reich *et al.* 2021). Milkweed samples were cleaned by rinsing with DDI and the use of an ultrasonic bath (~10 minutes), then dry ashed at 600 °C for 4 hours to assist digestion. Samples were digested in 16 M HNO_3_ (TraceMetal™ Grade; Fisher Chemical, Canada) for 16 hours at 120 °C. Ultraclean hydrogen peroxide was added to some samples to complete the digestion. After digestion, the samples were dried on a hot plate at 90 °C then re-dissolved in 1 mL of 2% v/v HNO_3_. One-tenth of the sample was separated out and diluted to 3 mL 2% v/v HNO_3_ for trace element analysis using the ICP-MS at the University of Ottawa.

Element concentrations were prepared by bracketing each set of samples with calibration standards prepared using single element certified standards purchased from SCP Science Inc. (Montreal, QC, Canada). Detection limits were conservatively estimated as three times the standard deviation of the blanks; 28 Cd and 5 Ni were below the calculated detection limit and were thus removed from further analysis. Some remaining Cd and Ni were near the detection limit; otherwise, procedural blanks were negligible (< 5%). Additionally, Mg were ~150 times higher than the calibration points and Cd were ~100 times lower than the calibration points; therefore, the absolute concentrations of these elements are uncertain, but the relative values are likely meaningful. Due to differences between the set-up of five analytical runs, the number of milkweed samples analyzed for each element differ (Table 2). Intra-run precision of standards was acceptable (RSD < 10%) for all elements except Mg (20%) and Al (15%). Between-run precision of peach leaves (n = 3; SRM 1547 Peach Leaves, National Institute of Standards and Technology) was found to be poor for all elements except Pb, Mn, and Al (Table S1). The sample concentrations are calculated based on the mass of the sample after it is dry ashed, a process which removes carbon, hydrogen, nitrogen, and some oxygen and volatile elements (e.g., Cl, Br, S) from the sample. Therefore, the milkweed concentrations cannot be directly compared to toxicological limits and the accuracy of the peach leaves standard measurements cannot be assessed. However, the relative concentrations are meaningful, allowing for the detection of relationships between milkweed and possible explanatory variables.

Of the 12 elements measured in the ashed milkweed leaves, Mg is the most abundant followed by Mn (Table S4). Milkweed leaves show broad ranges of elemental concentrations (Table S4). Strontium and Cd have the largest ranges with a ~70-fold difference between the minimum and maximum values; Mg has the smallest range with a substantial 5-fold difference. All elements show a positively skewed distribution.

#### 4.2 Elemental analysis of wild monarch wings

To estimate the natural exposure of wild monarch butterflies to metals in eastern North America, the wings of 100 monarch butterflies were tested for a suite of 12 elements (i.e., Al, Ba, Cd, Co, Cr, Cu, Mg, Mn, Ni, Pb, Sr, and Zn). The butterflies were captured in the spring of 2011 from sites in Texas, Oklahoma, and Missouri as part of a previous study (Flockhart *et al.* 2013). These butterflies are assumed to be members of the overwintering generation captured during their return migration from the overwintering sites in central Mexico. Therefore, the natal origins of these monarchs could be anywhere in the USA or Canada, although recent isotope-based geographic assignment using hydrogen and strontium isotopes found that most of these individuals likely originated in Texas (Reich *et al.* 2021).

To prepare the samples, a single forewing was dry-cleaned using pressurized nitrogen gas (~10 psi for 2 minutes) then digested in 16 M HNO_3_ (TraceMetal™ Grade; Fisher Chemical, Canada) for 16 hours at 120 °C. Ultraclean hydrogen peroxide was added to some samples to complete the digestion. After digestion, the samples were dried on a hot plate at 90 °C then re-dissolved in 1 mL of 2% v/v HNO_3_. One-tenth of the sample was separated out and diluted to 3 mL 2% v/v HNO_3_ for trace element analysis using the ICP-MS at the University of Ottawa.

Element concentrations were prepared by bracketing each set of samples with calibration standards prepared using single element certified standards purchased from SCP Science Inc. (Montreal, QC, Canada). Detection limits were conservatively estimated as three times the standard deviation of the blanks; measurements of 3 Cd, 63 Cr, 1 Co, and 63 Ni measurements were below the calculated detection limit and were thus removed from further analysis. Some remaining Cd, Co, and Ni measurements were near the detection limit; otherwise, procedural blanks were negligible (< 5%). Additionally, Cd were ~100 times lower than the calibration points; therefore, the absolute values of Cd are uncertain, but the relative values are likely meaningful. Intra-run precision of standards was acceptable (RSD < 10%) for all elements except Mg and Cr, which should be interpreted with caution.

Magnesium was the most abundant element detected followed by Al (i.e., over 100 ng/mg). All other elements had mean concentrations below 20 ng/mg.

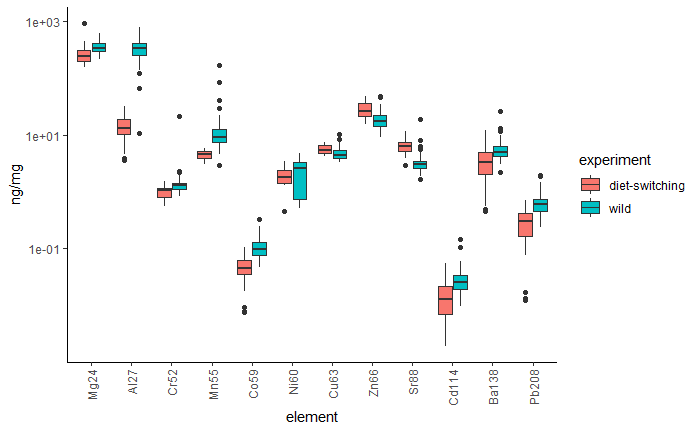

**Figure S5.** Boxplot comparing the concentrations of metals found in the forewings of wild monarch butterflies (blue; n = 100 except for Cr (n = 37), Co (n = 99), Ni (n = 37), and Cd (n = 97)) to those found in the diet-switching experiment (red; n = 18). Note the logarithmic scale on the y-axis. Wild monarchs tend to have higher metal concentrations than experimental monarchs, except for Cu, Zn, and Sr, which have lower concentrations.

#### 4.3 Comparison to permissible limits/toxicological studies

To test if the wild monarchs had metal concentrations that could indicate possible toxicological effects, the metal concentrations of the toxic metals As, Cd, Mn, Pb, and Tl measured in the monarch wings were compared to international permissible limits and toxicological studies on insects (Table S4). However, most of the studies looked at the concentrations of metals in food that produce adverse effects, rather than the concentrations of metals in insect tissue at which adverse effects occur. Regardless, examples of insect consumption thresholds are listed in Table S4. Few studies have tested the effects of metal exposure to monarch butterflies specifically, although notable exceptions have found negative impacts on monarch survival at doses of Zn as low as 344 ng/mg (Shephard *et al.* 2020) and high Na has been associated with decreased survival (Snell-Rood *et al.* 2014). However, a more recent study found no effects on monarch survival at Na concentrations of 2090 ng/mg and Zn of 555 ng/mg (Shephard *et al.* 2021). For Pb and Cd, international permissible limits as set by the World Health Organization (WHO) and the United Nation's Food and Agriculture Organization (FAO) for human food (FAO and WHO 2019) were used. Human-derived limits are non-protective for insects and that adverse physiological and behavioral effects can be detected well below set permissible limits (Hopkin 1990, Monchanin *et al.* 2021); thus, these limits should be considered to be overestimates. Few studies have investigated the toxicological effects on Mn on insects (Ben-Shahar 2018) but it is clear that effects are highly species-specific. For example, foraging behaviour of the European honeybee is impacted at Mn concentrations of 2750 ng/mg, the forest cockchafer (*Melolontha hippocastani*) exhibits reduced fitness at 1805 ng/mg, *Cabera pusaria* (L.) caterpillars were negatively affected at 695 ng/mg (Martinek *et al.* 2020), and *Lymantria dispar* (L.) moths were detected to have reduced survival at 4087 ng/mg (Kula *et al.* 2014). A study on *Spodoptera exigua* (Hübner) moths found minimum sublethal (i.e., larval weight loss) concentrations of Co, Ni, and Cu to be 15 ng/mg, 140 ng/mg, and 140 ng/mg, respectively (Cheruiyot *et al.* 2013).

| **Table S4.** Mean, standard deviation, and maximum concentration of the 12 metals measured in wild ashed milkweed and wild monarch butterflies across the eastern USA. Permissible limits of certain toxic elements are listed and occurrences above these limits are marked in bold text*.* | | | | | | |
| --- | --- | --- | --- | --- | --- | --- |
| metal | ashed milkweed (ng/mg) | | monarch (ng/mg; n=100) | | consumption threshold (ng/mg) | reference |
|  | mean ± SD (n) | range | mean ± SD | maximum |  |  |
| Al | 431±509 (85) | 71-3847 | 333±122 | 777 | 160 | (Kijak *et al.* 2014) |
| Cd | 0.11±0.10 (57) | 0.03-0.52 | 0.03±0.02 | **0.15** | 0.05-2 ­­­_­_ | (FAO and WHO 2019) |
| Cr | 1.84±1.07 (85) | 0.42-5.57 | 1.2±2.1 | 22 |  |  |
| Cu | 172±104 (85) | 51-472 | 4.7±1.3 | 10 | 140 | (Cheruiyot *et al.* 2013) |
| Mn | 880±980 (85) | 188-8114 | 12.9±18.0 | 167 | 695-4087 | (Kula *et al.* 2014, Søvik *et al.* 2015, Martinek *et al.* 2018, 2020) |
| Ni | 15.8±14.1 (80) | 1.4-74.1 | 0.9±1.3 | 4.8 | 140 | (Cheruiyot *et al.* 2013) |
| Pb | 1.38±1.01 (85) | 0.31-6.78 | **0.6±0.3** | **2** | 0.01-1 | (FAO and WHO 2019) |
| Zn | 460±299 (85) | 75-1688 | 19.1±6.9 | 48 | 344 | (Shephard *et al.* 2020) |
| Co | 0.96±1.03 (85) | 0.27-6.58 | 0.11±0.05 | 0.32 | 15 | (Cheruiyot *et al.* 2013) |
| Mg | 51932±17697 (85) | 26398-139666 | 354±86 | 618 |  |  |
| Ba | 309±286 (10) | 62-820 | 5.8±2.9 | 26 |  |  |
| Sr | 399±410 (85) | 37-2605 | 3.3±1.8 | 19 | NA | NA |

#### 4.4 Predictors of milkweed elemental composition

To identify factors that are potentially predictive of ashed milkweed element concentrations, the top-down linear mixed model selection approach suggested by Zuur et al. (2009) was followed for each element, excluding Ba due to low sample size. Previous studies have found that element concentrations in the soil and degree of urbanization can strongly influence milkweed elemental concentrations (Mitchell *et al.* 2020). Therefore, spatial geochemical soil data from three soil horizons (i.e., top 0-5 cm, A horizon, and C horizon) were downloaded from the USGS (Smith *et al.* 2013). Point data were interpolated for each of the elements and soil horizons using inverse distance weighting using ArcGIS Pro (Esri, Redlands, California, USA). Element concentrations in the three soil horizons were highly correlated, so the A horizon was chosen as a potential explanatory variable as this horizon is most likely to be accessed by the roots of milkweed and likely integrates both residual and transported (e.g., aeolian, anthropogenic) soils. Additionally, land use classifications from the National Land Cover Database (NLCD) were gathered (Yang *et al.* 2018, Jin *et al.* 2019, Homer *et al.* 2020, Dewitz and U.S. Geological Survey 2021). Classifications were simplified into four categories: natural, agricultural, wetland, and urban. Soil element concentrations and land use classifications were extracted from the spatial surfaces at the location of the milkweed collection sites. Linear mixed models were fit using the *nlme* package (Pinheiro *et al.* 2020) in R (R Core Team 2013) and final models were selected using AICs. Fixed effects included land use, soil elemental concentration and their interaction; random effects included ICP-MS analytical run, sampling year, sampling month, and milkweed species. Models including spatial autocorrelation structures (gaussian, spherical, and exponential) were also compared. Distributions of the response (i.e., ashed milkweed elemental concentration) and explanatory variables were inspected for normality and all milkweed elements and the soil concentrations of some elements (i.e., Cd, Cu, Co, Ni, Pb) were natural log transformed.

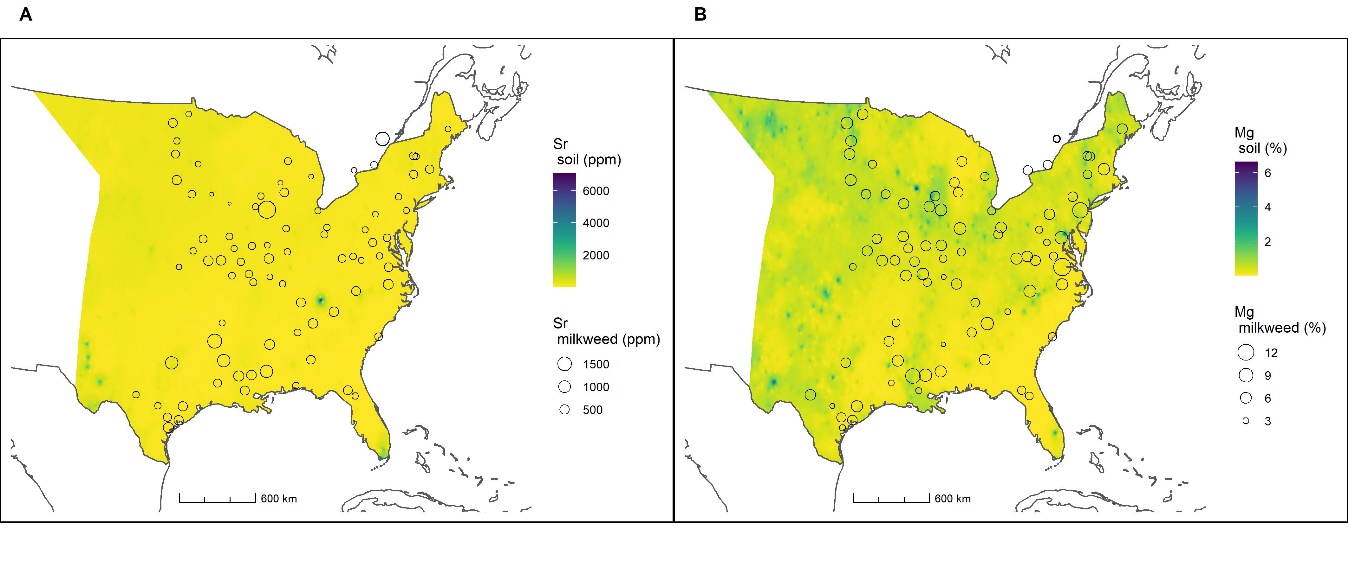

**Figure S6.** Maps demonstrating the weak correlations between metal concentrations in ashed milkweed samples and soil in the eastern USA. Metal concentrations in soil were interpolated from USGS soil samples (A horizon) (Smith *et al.* 2013). **(A)** Strontium concentration in the soil and milkweed are weakly correlated. The concentration in soil was not highlighted as an important variable by the model selection process (Table S5). **(B)** Likewise, Mg concentrations in the soil and the milkweed are weakly correlated and Mg concentration in the soil was not selected as an important predictor by the model selection procedure (Table S5).

Final models of ashed milkweed element concentrations show high variability in the explanatory variables selected for each element (Table S5). All final models except those for Mg and Sr include concentration in soil as an explanatory variable; however, this effect is only significant for Ni and Al. There is a negative relationship between the concentration of Cu and Cd in soil and in milkweed (Table S5); the concentration in milkweed of all other elements increases with higher concentration in soil. Landuse is only implicated as a relevant factor for Mg, Sr, Cd and Pb; this influence is mainly driven by the wetland category. A temporal variable (i.e., capture month or year) is included in the final model of Mn, Ni, Pb, Zn, Mg, and Sr. Overall, the linear mixed models have poor goodness-of-fit and exhibit considerable model uncertainty via intermediate models with similar AICs.

| **Table S5.** Final models for the concentration of each metal in ashed milkweed following the model selection procedure outlined in Zuur et al. (2009). Dashes indicate that the variable was not selected for the final model. All response variables and the soil concentrations of Cd, Cu, Co, Ni, and Pb were natural log-transformed to meet the assumption of normality. Pseudo R^2^ was calculated based on Nakagawa *et al.* (2017). *p < 0.05 | | | | | | | | | | |
| --- | --- | --- | --- | --- | --- | --- | --- | --- | --- | --- |
| metal | model fit | | fixed effect | | | random effect | | | | spatial autocorrelation  structure |
|  | AIC | Pseudo R² | soil (β±SE, df, t) | landuse (Wald test; df, F) | soil * landuse (Wald test; df, F) | species  (SD) | analytical run (SD) | Year | Month |  |
| Al | 212 | 0.30 | 0.13±0.07, 72, 2.0* | - | - | 0.42 | - | - | - | - |
| Cd | 127 | - | -0.71±0.47, 55, -1.5 | 3,0.4 | 3, 3.6* | - | - |  |  | gaussian |
| Cr | 127 | 0.51 | 0.0025±0.0014, 79, 1.8 | - | - | - | 0.43 |  |  | - |
| Cu | 92 | - | -.065±0.080, 61, -0.8 | - | - | 0.053 | 0.51 | - | - | - |
| Mn | 172 | - | 0.00015±0.00017, 64, 0.9 | - | - | 0.63 | - | - | 0.0005 | exponential |
| Ni | 176 | - | 0.25±0.10, 48, 2.4* | - | - | 0.73 | 0.34 | - | 0.31 | - |
| Pb | 165 | 0.32 | 0.59±0.42, 72, 1.4 | 3, 5.9* | 3, 2.8* | - | - | - | 0.20 | - |
| Zn | 142 | - | 0.0015±0.0021, 66, 0.72 | - | - | 0.66 | - | 0.00004 | - | - |
| Co | 177 | 0.42 | 0.075±0.078, 79, 1.0 |  |  | - | 0.53 | - | - | gaussian |
| Mg | 48 | - | - | 3, 1.72 | - | 0.22 | - | 0.00002 | 0.000003 | - |
| Sr | 191 | - | - | 3, 4.1* | - | 0.39 | - | 0.44 | - | exponential |

We expected to find strong relationships between the element concentrations in soil and milkweed; strong relationships would suggest that current maps of metal concentrations in soil (USGS) could be used like an isoscape for geographic assignment (i.e., a “metalscape”). However, we found weak and non-significant relationships between the soil and biosphere for 7 of the 11 elements (e.g., Figure S6). Both Sr and Mg were suggested as geolocation tools that should be further explored (main text, Table 1), but concentration in soil was not chosen by the model selection procedure as an important predictor for either of these metals (Table S5 and Figure S6). This suggests that more research is needed to understand the spatial patterns of metals in insect tissue before single metals can be used for geolocation purposes. Previous studies have also found similarly weak correlations in both natural (Wieringa et al., 2020) and controlled laboratory (Lin et al., 2021) conditions. Weak correlations are likely influenced by the fact that the elemental soil concentrations used in this study are interpolated from samples of the total soil and may not reflect the bioavailable component of soil elements accessible to the plants (Tibbett *et al.* 2021). Soil samples that are processed to represent the bioavailable portion of soil elements may yield stronger relationships (Peijnenburg and Jager 2003). Many of the final linear models included milkweed species as a random effect and indicate substantial differences between species (Table S5). The inclusion of this term suggests that there may be inter-specific differences in metal uptake, regulatory, or storage mechanisms even among milkweed species. Alternatively, milkweed species are known to occupy distinct ecosystems (Pocius *et al.* 2018) and thus this variable could be selected by the model as a proxy for ecological and environmental conditions at a higher resolution than the landuse variable. The poor fit of the linear mixed models indicates that explanatory variables not included in the full model are likely important in determining the concentration of elements in milkweed. Elemental concentrations in milkweed may also be influenced by factors such as distance to anthropogenic sources of metals (Fritsch *et al.* 2011), developmental stage of the plant, and soil type. Nevertheless, previous studies have found that the weak relationships found between single elements does not necessarily preclude the geolocation ability of trace elements when combined into a chemoprint (Wieringa *et al.* 2020).
